# Neuroanatomical changes observed over the course of a human pregnancy

**DOI:** 10.1101/2023.12.14.571688

**Authors:** Laura Pritschet, Caitlin M. Taylor, Daniela Cossio, Joshua Faskowitz, Tyler Santander, Daniel A. Handwerker, Hannah Grotzinger, Evan Layher, Elizabeth R. Chrastil, Emily G. Jacobs

**Affiliations:** Department of Psychological & Brain Sciences, University of California, Santa Barbara, CA; Department of Neurobiology and Behavior, University of California, Irvine, CA; Section on Functional Imaging Methods, Laboratory of Brain and Cognition, National Institute of Mental Health, National Institutes of Health, Bethesda, MD; Neuroscience Research Institute, University of California, Santa Barbara, CA

**Keywords:** pregnancy, brain structure, MRI, sex steroid hormones, precision imaging, women’s health

## Abstract

Pregnancy is a period of profound hormonal and physiological change experienced by millions of women annually, yet the neural changes unfolding in the maternal brain throughout gestation have not been studied in humans. Leveraging precision imaging, we mapped neuroanatomical changes in an individual from preconception through two years postpartum. Pronounced decreases in gray matter volume and cortical thickness were evident across the brain, which stand in contrast to increases in white matter microstructural integrity, ventricle volume, and cerebrospinal fluid, with few regions untouched by the transition to motherhood. This dataset serves as the first comprehensive map of the human brain across gestation, providing an open-access resource for the brain imaging community to stimulate further exploration and discovery.

---

Worldwide, nearly 85% of women experience one or more pregnancies in their lifetime (CDC, 2023), with 140 million women becoming pregnant each year. Over an approximately 40-week gestational window the maternal body undergoes profound physiological adaptations to support the development of the fetus, including increases in plasma volume, metabolic rate, oxygen consumption, and immune regulation (Thornburg et al., 2015). These rapid adaptations are initiated by hundred-to thousand-fold increases in hormone production, including estrogen and progesterone. These neuromodulatory hormones also drive significant reorganization of the central nervous system. Evidence from animal models and human studies converge on pregnancy as a period of remarkable neuroplasticity (Brunton & Russell et al., 2008; Haim et al., 2017; Barrière et al., 2021; Celik et al., 2022; Puri et al., 2023; Chaker et al., 2023; see Diamond et al., 1971 for one of the earliest known observations). Gestational increases in steroid hormone synthesis drive neurogenesis, dendritic spine growth, microglial proliferation, myelination, and astrocyte remodeling (for review, see Servin-Barthet et al., 2023). These cellular changes are pronounced in brain circuits that promote maternal behavior. For example, Ammari and colleagues (2023) recently discovered steroid hormones’ ability to fine-tune the response properties of galanin neurons in the rodent medial preoptic area of the hypothalamus (mPOA); changes that enhance dams’ sensitivity to sensory cues from newborn pups.

In humans, reductions in gray matter volume (GMV) have been observed postpartum (Hoekzema et al., 2017, 2022; Martínez-García, et al., 2021a; Spalek et al., 2024), particularly in regions central to theory-of-mind processing (Hoekzema et al., 2017). These GMV changes persist at six years postpartum (Martínez-García et al., 2021b) and are traceable decades later (De Lange et al., 2019; Orchard et al., 2020), underscoring the permanence of this major remodeling event. And yet the changes that occur within the maternal brain *during* gestation itself are virtually unknown. A recent study by Paternina-Die and colleagues (2024) offers intriguing clues. Women were scanned preconception, once in the 3^rd^ trimester, and again in the postpartum period, revealing a reduction of cortical volume observable in the late pregnancy scan. These findings suggest that pregnancy is a highly dynamic period for neural remodeling, yet neuroscientists lack a detailed map of how the human brain changes throughout the gestational period.

Here, we conducted the first precision imaging study of pregnancy in which a healthy 38-year-old primiparous woman underwent 26 MRI scans and venipuncture beginning 3 weeks pre-conception through two years postpartum. We observed widespread reductions in cortical GMV and cortical thickness (CT) occurring in step with advancing gestational week and the dramatic rise in sex hormone production. Remodeling was also evident within subcortical structures, including the ventral diencephalon, caudate, thalamus, putamen, and hippocampus. High-resolution imaging and segmentation of the medial temporal lobe extend these findings further, revealing specific volumetric reductions within hippocampal subfields CA1, CA2/3, and parahippocampal cortex. In contrast to widespread decreases in cortical and subcortical GMV, correlational tractography analyses revealed non-linear increases in white matter quantitative anisotropy (QA) throughout the brain —indicating greater tract integrity— as gestational week progressed. Together, these findings are the first to reveal the highly dynamic changes that unfold in a human brain across pregnancy, demonstrating a capacity for extensive neural remodeling well into adulthood.

## Results

### Serological evaluations

Serological evaluations captured canonical hormone fluctuations characteristic of the prenatal, perinatal, and postnatal periods (**Fig. 1B**). Serum hormone concentrations increased significantly over the course of pregnancy and dropped precipitously postpartum (*pre-conception*: estradiol (E) = 3.42 pg/mL, progesterone (P) = 0.84 ng/mL; *3 weeks prior to parturition*: E = 12,400 pg/mL, P = 103 ng/mL; *3 months after parturition*: E = 11.50 pg/mL, P = 0.04 ng/mL).

**Figure 1.**
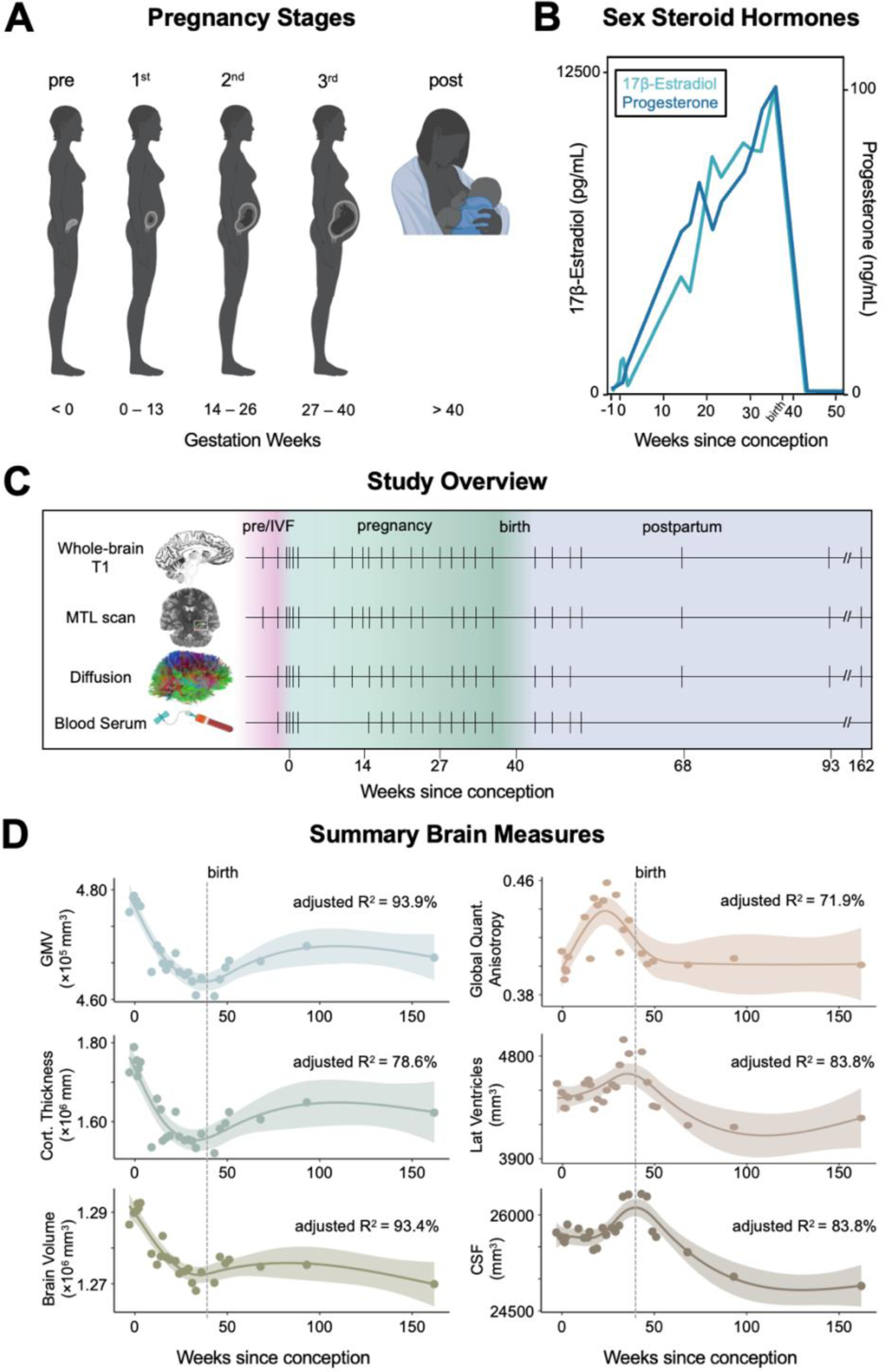
Precision imaging reveals neuroanatomical changes throughout gestation. **A)** Standard medical demarcations for pregnancy stages imesters) by gestation week (created with BioRender.com). **B)** Steroid hormones increased significantly over the course of pregnancy opped precipitously postpartum, as is characteristic of the pre- and postnatal periods. **C)** A healthy 38-year-old primiparous woman ent 26 scanning sessions from 3 weeks preconception through two years postpartum. Scans were distributed throughout ception (4 scans), first trimester (4 scans), second trimester (6 scans), third trimester (5 scans), and postpartum (7 scans); tick marks e when major measures were collected, colors denote pregnancy stage. The participant underwent in-vitro fertilization (IVF) to e pregnancy, allowing for precise mapping of ovulation, conception, and gestation week. **D)** Summary (i.e., total) brain measures e course of the experiment. Gray matter volume, cortical thickness, and total brain volume decreased over the course of pregnancy *ethods*), with a slight recovery postpartum. Global quantitative anisotropy, lateral ventricle and cerebrospinal fluid volumes displayed ear increases across gestation, with a notable rise in the second and third trimesters before dropping sharply postpartum. Shaded s represent a 95% confidence interval derived from generalized additive models; dashed line indicates parturition. *Abbreviations*: in-vitro fertilization; MTL = medial temporal lobe; GMV = gray matter volume; CSF = cerebrospinal fluid.

### Whole-brain dynamics from baseline through postpartum

To begin, we characterized broad neuroanatomical changes over the course of the entire experimental window (baseline – 2 years postpartum; 26 scans; **Fig. 1D**). Generalized additive models revealed strong non-linear (effective degrees of freedom > 3) relationships between weeks since conception and summary brain metrics. Total GMV (*F =* 27.87, *p <* .001*, deviance explained* = 93.9%, *R^2^_adj_ = 0.91*), summary CT (*F =* 15.79*, p <* .001*, deviance explained* = 78.6%, *R^2^ = 0.*75) and total brain volume (*F =* 26.12*, p <* .001*, deviance explained* = 93.4%, *R^2^_adj_ = 0.*90) linearly decreased during gestation and appeared to partially rebound postpartum. In contrast, global microstructural integrity (QA) of white matter increased throughout the first *QA, F =* 6.80*, p <* .002*, deviance explained* = 71.9%, *R^2^ = 0.*63). We also observed non-linear patterns of lateral ventricle expansion *(F =* 10.44*, p <* .001*, deviance explained* = 83.8%, *R^2^ = 0.*77) and increased cerebrospinal fluid (CSF; *F =* 13.32*, p <* .001*, deviance explained* = 83.8%, *R^2^ = 0.*79) rising in the second and third trimesters before dropping sharply postpartum.

### Cortical volume and thickness changes tied to gestation

We then narrowed the aperture to capture changes unfolding within gestation itself (baseline – 36 weeks pregnant; 19 scans). Relationships between summary brain metrics were evident over the gestational period: total brain volume, GMV, and CT were positively associated with one another, whereas CSF and global QA demonstrated negative relationships with GMV (**Fig. S1**).

Changes in GMV were near-ubiquitous across the cortical mantle (**Fig. 2A**). Most large-scale brain networks exhibited decreases in GMV (**Fig. 2B, Table S1**); indeed, 80% of the 400 regions of interest (ROIs) demonstrated negative relationships between GMV and gestation week (**Fig. 2A**, **Table S2**). Together, these results provide evidence of a global decrease in cortical volume across pregnancy. Several sensory and attention subnetworks were particularly sensitive to gestation, including the Control (B), Salience/Ventral Attention (A), Dorsal Attention (B), Default (A), and Somatomotor (A,B) networks (**Table S1**). Regions driving these network-level changes include the bilateral inferior parietal lobe, post central gyri, insulae, prefrontal cortex, posterior cingulate, and somatosensory cortex (**Fig. 2C**, **Table S2**; see **Tables S1 and S3–4** for validation of findings using alternate pipeline). These regions and associated brain networks appear to decrease in volume at a faster rate than the rest of the brain throughout pregnancy, as determined by a subsequent analysis controlling for total GMV (**Tables S1–2**). GMV reductions were also significantly correlated with the participant’s estradiol and progesterone concentrations (**Table S1**). A highly similar pattern of results was observed when examining pregnancy-related cortical thickness changes (**Fig. S3, Tables S4–5**). Significant reductions in cortical GMV over gestation remained after controlling for standard quality control metrics, albeit with some influence on the magnitude and location of the observed effects (see **Figures S4–5**).

**Figure 2.**
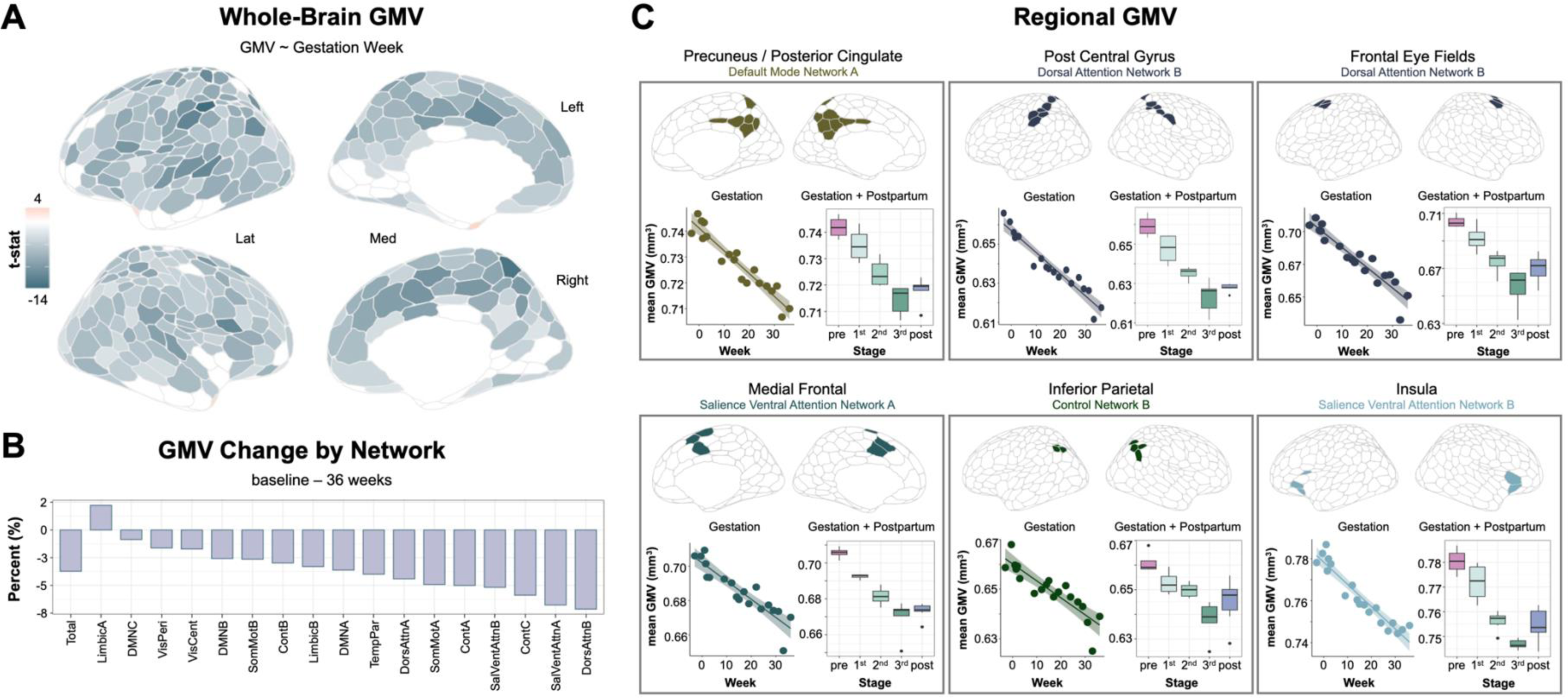
Cortical gray matter volume showed widespread change through gestation and postpartum. **A)** Multivariate regression analyses reveal largely negative relationships between gestation week and regional GMV, with only a minority of regions unaffected or increasing over the gestational window (baseline – 36 weeks). All associations presented here were corrected for multiple comparisons (FDR at *q* < 0.05; non-significant test statistics were set to zero for interpretability). **B)** Average network change was calculated by estimating GMV percent change from baseline (initial) to 36 weeks gestation (final). Attention and Control Networks appear most affected. **C)** Six representative regions, classified by major subnetworks, that exhibit pronounced GMV change across gestation. For each panel, we display a scatterplot between average GMV of the ROIs and gestation week (left; gestation sessions only), and summary ROI GMV by pregnancy stage across the whole study (right; gestation and postpartum sessions). All statistical tests were corrected for multiple comparisons (FDR at *q* < 0.05). Color-coded by network affiliation (see **Fig. S2**). N.b., shown here are raw data values (see **Tables S1–2** and **Supplementary File 2** for exhaustive list). *Abbreviations*: GMV = gray matter volume; Lat = lateral; Med = medial; DMN = Default Mode Network; VisPeri = Visual Peripheral Network; SomMot = Somatomotor Network; VisCent = Visual Central Network; Cont = Control Network; TempPar = Temporal Parietal Network; DorsAttn = Dorsal Attention Network; SalVentAttn = Salience / Ventral Attention Network. Brain visualizations created with R-package *ggseg* (Mowinckel and Vidal-Piñeiro, 2020).

In contrast, GMV within regions of the Default Mode (C), Limbic (A,B), and Visual Peripheral Networks buck the global trend by slightly increasing (e.g., temporal poles), remaining constant (e.g., orbitofrontal cortex), or reducing at a much slower rate (e.g., extrastriate cortex) than total GMV (**Fig. 2A–B, Tables S1–2**). Cortical thickness changes in these regions exhibit similar patterns (**Fig. S3, Tables S4–5**).

### Subcortical gray matter volume changes tied to gestation

Consistent with the broader cortical reductions in GMV, several subcortical regions significantly reduced in volume across gestation (**Fig. 3A**, *left*). This included bilateral ventral diencephalon (right hemisphere values shown in **Fig. 3A**, *right*; encompasses hypothalamus, substantia nigra, mammillary body, lateral geniculate nucleus, and red nucleus among others; Makris et al., 2008), caudate, hippocampus, and thalamus, along with left putamen and brain stem (**Table S6**, *q* < 0.05).

**Figure 3.**
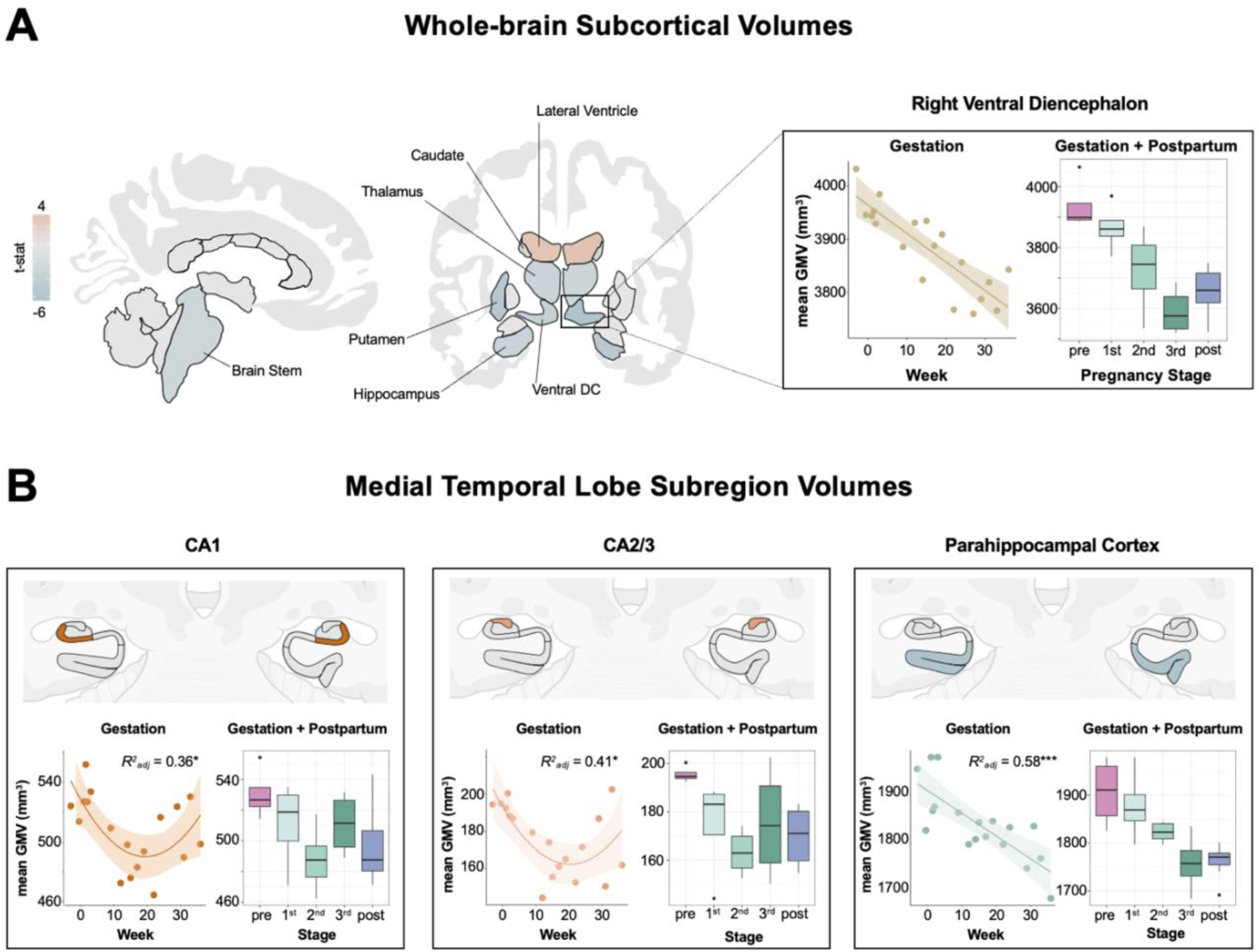
Subcortical gray matter volume changed throughout gestation. **A)** Multivariate regression analyses revealed largely negative relationships between gestation week and subcortical GMV regions over pregnancy, including bilateral thalamus, caudate, hippocampus, ventral diencephalon (encompassing hypothalamus, substantia nigra, mammillary body, and red nucleus), and left caudate. Lateral ventricles displayed the only positive relationships with gestation week (also depicted in Fig. 1D). The whole-brain subcortical GMV estimates shown here were derived via FreeSurfer and ‘aseg’ subcortical segmentation. FDR-corrected at *q* < 0.05. *Inset:* Right ventral diencephalon displayed the strongest negative association with gestation (left) and did not return to baseline postpartum (right). **B)** The participant’s hippocampus and surrounding cortex were segmented into seven bilateral subregions. CA1, CA2/3 and parahippocampal cortex were negatively associated with gestation week and did not return to baseline postpartum. FDR-corrected at *q* < 0.05; ***q < 0.001; *q < 0.05. For both panels, non-significant regions were set to zero (gray) for interpretability. See **Fig. S6** for complete labelling of regions in both segmentations. Abbreviation: DC = diencephalon; GMV = gray matter volume. Brain visualizations created with R-package *ggseg* (Mowinckel and Vidal-Piñeiro, 2020).

Next, high-resolution segmentation of the medial temporal lobe allowed us to interrogate subcortical structures at a finer resolution, revealing non-linear volumetric decreases in CA1 and CA2/3 (*CA1: F(2,15) =* 5.84, *q* = .031, *R^2^_Adj_ =* 0.36; *CA2/3: F(2,15)=* 6.82, *q* = .027, *R^2^_Adj_ =* 0.41) across gestation (**Fig. 3B**, *left and middle*). Parahippocampal cortex exhibited linear volumetric decreases across gestation (PHC; *F(1,16) =* 24.87*, q <* .001*, R^2^_Adj_ =* 0.58; **Fig. 3B**, *right*) which was also tied to estradiol (*F(1,12) =* 20.21*, q =* .005*, R^2^_Adj_ =* 0.60). All three relationships remained significant after proportional correction for total GMV. There was no significant change in other subregions or total volume of the hippocampal body, or in the parahippocampal gyrus (**Table S7**, **Fig. S8**).

### White matter microstructure changes tied to gestation

In contrast to decreasing global GMV, correlational tractography of white matter, which tests for linear trends in the data, revealed increasing microstructural integrity across the whole brain during gestation (**Fig. 4A**), concomitant with the rise in 17ß-estradiol and progesterone (*q* < .001) (**Fig. S9**). Tracts displaying robust correlations with gestational week included the corpus callosum, arcuate fasciculus, inferior fronto-occipital fasciculus, and inferior longitudinal fasciculus (**Fig. 4B**), as well as the cingulum bundle, middle and superior longitudinal fasciculus, corticostriatal tracts, and corticopontine tracts (see **Table S9** for complete list).

**Figure 4.**
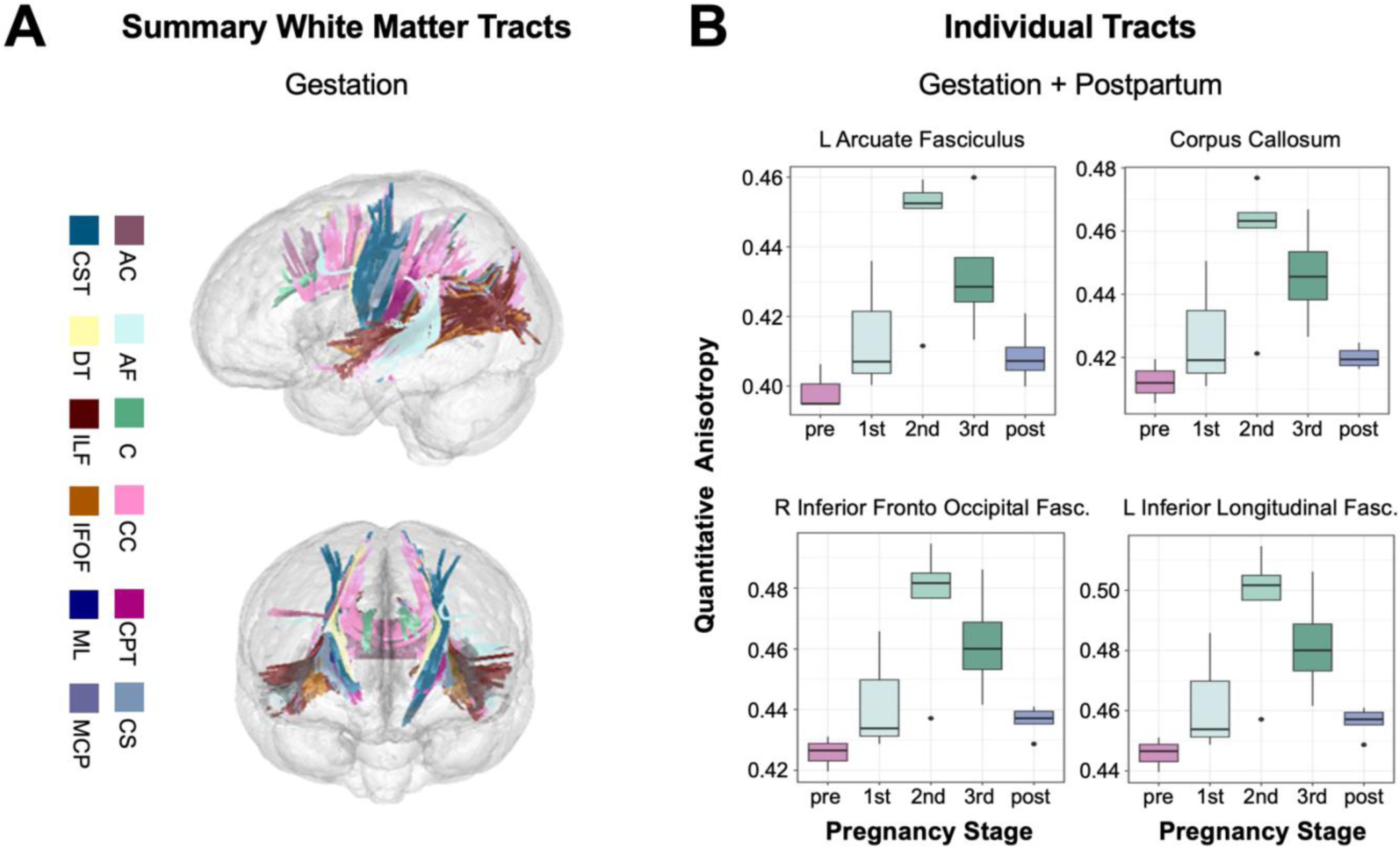
White matter microstructure changes over the course of the experiment. **A)** Numerous white matter tracts demonstrate increasing quantitative anisotropy (QA) in relation to advancing gestation week, as determined by correlational tractography analysis. **B)** Summary of QA values by pregnancy stage for representative ROIs significantly tied to gestation. Tractometry was used to extract quantitative anisotropy values. Abbreviations: AC = anterior commissure; AF = arcuate fasciculus; C = cingulum bundle; CC = corpus callosum; CPT = corticopontine tracts; CS = corticostriatal tracts; CST = corticospinal tracts; DT = dentothalamic tract; ILF = inferior longitudinal fasciculus; IFOF = inferior frontal occipital fasciculus; ML = medial lemniscus; MCP = middle cerebellar peduncle.

### Comparing brain changes across pregnancy against densely-sampled controls

We then compared the changes in GMV across gestation to that of typical variability over time, derived from eight densely-sampled controls (Filevich et al., 2017). The GMV changes we see across pregnancy far exceed normative brain variability (**Fig. S11**). On average, change in cortical GMV was nearly three times higher than controls scanned over a similar duration (**Fig. S11A–B).** This extends to MTL subfields, wherein change in volume was three-to four-times greater across gestation than normative brain variability (**Fig. S11C–D)**. We contextualized these findings further by comparing gestational GMV change against our participant’s pre-conception brain volumes; average GMV change during pregnancy was six times (cortical) and three times (MTL) higher than the variability observed between baseline sessions.

## Discussion

Converging evidence across mammalian species points to pregnancy as a remarkable period of neuroplasticity, revealing the brain’s ability to undergo adaptive, hormonally-driven neuroanatomical changes beyond adolescence (Dulac et al., 2014; Hoekzema et al., 2017, 2022; Carmona et al., 2019; Martínez-García, et al., 2021a; Pawluski et al., 2022; Paternina-Die et al., 2024). Investigations that compare women pre- and postpartum provide the strongest evidence to date that the human brain undergoes such neural changes (Servin-Barthet et al., 2023; Martínez-García et al., *in press*). But what about pregnancy itself? Over what time-course do anatomical changes in the maternal brain manifest? Are they tied to the substantial increase in sex hormone production? Here, we begin to address these outstanding questions. This paper and corresponding open-access dataset offer neuroscientists a detailed map of the human brain across gestation, a resource for which a wide-range of previously unattainable neurobiological questions can now be explored.

Our findings from this precision imaging study show that pregnancy is characterized by sweeping reductions in gray matter volume, cortical thinning, and enhanced white matter microstructural integrity that unfold week by week. These changes were also tied to the significant rise in steroid hormone concentrations over pregnancy. Some of these changes persist at two years postpartum (e.g., global reductions in GMV, CT), while others, including markers of white matter integrity, appear to be transient. Ventricular expansion and contraction parallel these cortical changes. These widespread patterns, and the notable increase in CSF volume across gestation, could reflect increased water retention and subsequent compression of cortical tissue. However, the persistence of these changes at two years postpartum and regional variation in GMV, CT, and QA, hint at cellular underpinnings, such as alterations in glia or neuron number, synaptic density, and myelination. Future studies of the relationship between fluid dynamics and volumetric changes will help clarify the factors that drive global neural changes during pregnancy; such insights will have broad implications for maternal health (e.g., neurological effects tied to pre-eclampsia or edema).

Critically, dynamic neural changes occurred *within* the pregnancy window itself, a nuance not captured by studies limited to pre-versus post-pregnancy comparisons. For example, we observed large increases in white matter microstructural integrity (QA) throughout the first and second trimesters of pregnancy, but these measures fully returned to baseline values by the first postpartum scan. This pattern may explain why previous studies report no pregnancy-related differences in white matter tractography (Hoekzema et al., 2022). Other measures, such as GMV and CT, decreased throughout gestation and displayed only a modest rebound postpartum. These non-linear patterns suggest that only quantifying pre- and postpartum brain structure may overlook the full range of changes that unfold within the gestational window — and underrepresent the brain’s metamorphosis during pregnancy. Further, although observed changes were largely global, some regions displayed notable stability (e.g., extrastriate cortex). The subcortical region that displayed the strongest relationship with gestation week was the ventral diencephalon, which encompasses the hypothalamus and subsequent medial preoptic area and paraventricular nucleus — structures critical for inducing maternal behavior (Ammari et al., 2023 and Spalek et al., 2024). The hippocampus exhibited a reduction in volume across gestation, and with higher spatial resolution, this reduction was revealed to be driven by changes in CA1 and CA2/3 subfield volumes, while other hippocampal subfields remained stable. Adjacent parahippocampal cortex within the medial temporal lobe also exhibited volume reduction across gestation. While our hippocampal findings are consistent with pre/post studies of pregnancy (Hoekzema et al., 2017), the precision lens applied within gestation revealed the non-linear nature of this reduction. Recapitulating and clarifying these regionally specific patterns of volume change throughout the medial temporal lobe merits further investigation.

Similar precision imaging studies have captured dynamic brain reorganization across other neuroendocrine transitions, such as the menstrual cycle (see review: Pritschet et al., 2021), underscoring the powerful role steroid hormones play in shaping the mammalian brain (Taxier et al., 2020). Endocrine changes across pregnancy dwarf those that occur across the menstrual cycle, which highlights the critical need to map the brain’s response to this unique hormonal state. Broad physiological changes occur in tandem with the rise in steroid hormones, including changes in body mass composition, water retention, immune function, and sleep patterns (Servin-Barthet et al., 2023). These factors could be playing a role in the brain changes observed here, with some driving neurobiological changes and others, like water retention, potentially affecting MRI-based measurements. Note that, although cortical reductions in GMV over gestation were stable across analyses, accounting for certain quality control measures influenced the magnitude and location of these results. These metrics all fell within the standard range, but there may be meaningful reductions in signal that accompany volumetric reductions (e.g., increased CSF, decreased GM) — a methodological nuance that goes beyond the scope of this resource paper. Ultimately, identifying the shared and unique contributions of these factors to the neuroanatomical changes that unfold across gestation warrants further investigation. Deeply phenotyping a large and diverse cohort of women across pregnancy will open up new avenues of exploration, e.g., allowing researchers to link blood-based proteomic signatures to pregnancy outcomes; deploying wearable devices to monitor changes in sleep, cognition, and mood; and probing the broader social and environmental determinants of maternal health (Martínez-García et al., in press).

The neuroanatomical changes that unfold during matrescence have broad implications for understanding individual differences in parental behavior (Dulac et al., 2014; Hoekzema et al., 2017; Kohl et al., 2018; Rodrigo et al., 2016), vulnerability to mental health disorders (Pawluski et al., 2017; Barba-Müller et al., 2019) and patterns of brain aging (de Lange et al., 2019; Barth & de Lange, 2020; Orchard et al., 2020, 2023; Duarte-Guterman et al., 2023). Decreases in GMV may reflect “fine-tuning” of the brain by neuromodulatory hormones in preparation for parenthood (Pawluski et al., 2022). For example, in rodents, steroid hormones promote parental behavior by remodeling specific neural circuits in the medial preoptic area of the hypothalamus. These behavioral adaptations are critical to the dam’s ability to meet the demands of caring for the offspring (Ammari et al., 2023). Human studies have revealed GMV reductions in areas of the brain important for social cognition and the magnitude of these changes corresponds with increased parental attachment (Hoekzema et al., 2017). Deeper examination of cellular and systems-level mechanisms will improve our understanding of how pregnancy remodels specific circuits to promote maternal behavior.

Although studied to a lesser degree, ties between maternal behavior and white matter microstructure —particularly connectivity between temporal and occipital lobes— have been noted (Rodrigo et al., 2016). Here, we reveal pronounced GMV changes in regions within sensory, attention, and default mode networks over the gestational window. In parallel, we observed increased anisotropy in white matter tracts that facilitate communication between emotional and visual processing hubs (Herbert et al., 2018, Wang et al., 2018; Zekelman et al., 2022), including the inferior longitudinal fasciculus and inferior fronto-occipital fasciculus. Pinpointing the synchrony of gray and white matter changes that unfold in the maternal brain could be key to understanding the behavioral adaptions that emerge during and after pregnancy, such as honing the brain’s visual and auditory responses to infant cues and eliciting maternal behavior. Research into other major transition periods supports this idea. For instance, adolescence is a dynamic period characterized by region-specific, non-linear decreases in GMV and increases in WMV, maturational brain changes that are tied to gains in executive function and social cognition (Blakemore & Choudhury, 2006). For both adolescence (Blakemore and Dahl, 2010) and matrescence, the considerable rise in steroid hormone production appears to remodel the brain (see Carmona et al., 2019 for comparative analysis), promoting a suite of behaviors adaptive to that life stage. How specific neural changes give rise to specific behavioral adaptations has yet to be fully explored with respect to human pregnancy.

This precision imaging study mapped neuroanatomical changes across pregnancy in a single individual, precluding our ability to generalize to the broader population. To benchmark our findings, we compared the magnitude of GMV changes observed throughout pregnancy against data from non-pregnant individuals sampled over a similar time course. Doing so provided compelling evidence that pregnancy-related neuroanatomical shifts far exceed normative day-to-day brain variability and measurement error. Evidence suggests that white matter microstructure remains fairly stable over a six-month period (Lövdén et al., 2010), but more studies are needed to compare the degree of white matter changes observed during pregnancy to normative change over time. Further, sampling larger cohorts of women will generate much-needed normative models of brain change (akin to Bethlehem et al., 2022) throughout pregnancy to establish what constitutes a typical degree of neuroanatomical change expected during gestation and postpartum recovery.

These findings provide critical rationale for conducting further precision imaging studies of pregnancy in demographically enriched cohorts to determine the universality and idiosyncrasy of these adaptations and their role in maternal health. Are the changes observed in our participant reflective of the broader population? Do deviations from the norm lead to maladaptive outcomes? A precision imaging approach can help determine whether the pace of pregnancy-induced neuroanatomical changes drives divergent brain health outcomes in women, as may be the case during other rapid periods of brain development (Tooley et al. 2021). One in five women experiences perinatal depression (Wang et al., 2021) and while the first FDA-approved treatment is now available (Deligiannidis et al., 2023), early detection remains elusive. Precision imaging studies could offer clues about an individual’s risk for or resilience to depression prior to symptom onset, helping clinicians better determine when and how to intervene. Neuroscientists and clinicians also lack tools to facilitate detection and treatment of neurological disorders that co-occur, worsen, or remit with pregnancy, such as epilepsy, headaches, multiple sclerosis, and intracranial hypertension (Shehata et al., 2004). This new area of study—precision mapping of the maternal brain—lays the groundwork for a greater understanding of the subtle and sweeping structural, functional, behavioral, and clinical changes that unfold across pregnancy. Such pursuits will advance our basic understanding of the human brain and its remarkable ability to undergo protracted plasticity in adulthood.

## Materials and Methods (Online only)

### Participant

Our participant (author E.R.C.) was a healthy 38-year-old primiparous woman who underwent in-vitro fertilization (IVF) to achieve pregnancy. Previous studies reported no observable differences in neural changes pre-to post-pregnancy between women who conceived naturally versus women who conceived via IVF (Hoekzema et al., 2017), and doing so provides a controlled way of monitoring pregnancy status. The participant experienced no pregnancy complications (e.g., gestational diabetes, hypertension), delivered at full term via vaginal birth, nursed through 16 months postpartum, and had no history of neuropsychiatric diagnosis, endocrine disorders, prior head trauma, or history of smoking. The participant gave written informed consent and the study was approved by the University of California, Irvine Human Subjects Committee.

### Study Design

The participant underwent 26 magnetic resonance imaging (MRI) scanning sessions from 3 weeks prior to conception through two years postpartum (162 weeks), during which high-resolution anatomical and diffusion spectrum imaging scans of the brain were acquired. Scans were distributed throughout this period, including pre-pregnancy (4 scans), first trimester (4 scans), second trimester (6 scans), third trimester (5 scans), and postpartum (7 scans) (**Fig. 1C**). The first 6 sessions took place at the UCSB Brain Imaging Center (BIC), the final 20 sessions took place at the UCI Facility for Imaging and Brain Research (FIBRE). The majority of scans took place between 9am-2pm, limiting significant AM/PM fluctuations. The MRI protocol, scanner (Siemens 3T Prisma), and software (version MR E11) were identical across sites. Each scanner underwent weekly quality control checks for the duration of the study and passed all reports indicating no significant alterations in the geometry. To ensure the robustness of the findings, after the final study session the subject completed back-to-back validation scans at UCI and UCSB within a 12-hour window to assess reliability between scanners. Intraclass correlation coefficients (two-way, random effects, absolute agreement, single rater) reveal ‘excellent’ test-retest reliability between scanners, including: ROI-level GMV (*ICC* = 0.97, *95% CI*: 0.80–0.99); ROI-level CT (*ICC* = 0.96, *95% CI*: 0.90–0.98); MTL subfield volume (*ICC* = 0.99, *95% CI:* 0.97–0.99); and ROI-level QA (*ICC* = 0.94, *95% CI:* 0.91–0.97). Further, when examining the relationship between gestation week and GMV among UCI-only gestational sessions, findings were consistent (**Fig. S12**). Scanning a control participant over the same time course was not permissible due to the ongoing COVID-19 pandemic; however, the above quality checks suggest that within- and between-scanner variability is highly unlikely to have contributed meaningfully to the observed effects.

To monitor state-dependent mood and lifestyle measures, the following scales were administered on each experiment day: Perceived Stress Scale (PSS; Cohen et al., 1983), Pittsburgh Sleep Quality Index (PSQI; Buysse et al., 1989), State-Trait Anxiety Inventory for Adults (STAI; Spielberger and Vagg, 1984), and Profile of Mood States (POMS; Pollock et al., 1979). Correlation analyses between state-dependent measures, summary brain metrics, and gestation week revealed little to no relationships. The only exception to this was a moderate negative association between global QA and state anxiety (*spearman’s rho* = -0.59, *q* = 0.04; baseline – 36 weeks, n = 19). By making this data openly accessible, we encourage a more nuanced approach towards exploring mood and lifestyle measures in relation to brain changes over pregnancy.

### Endocrine Procedures

The participant underwent a blood draw (n = 19, **Fig. 1C**) prior to MRI scanning. Sex steroid concentrations were determined via ultra-sensitive liquid chromatography–mass spectrometry (LC-MS) at the Brigham and Women’s Hospital Research Assay Core (BRAC). Assay sensitivities, dynamic range, and intra-assay coefficients of variation were as follows: estradiol: 1.0 pg/ml, 1–500 pg/ml, <5% relative standard deviation (RSD); progesterone: 0.05 ng/ml, 0.05– 10 ng/ml, 9.33% RSD. Serological samples were not acquired in five sessions due to scheduling conflicts with UC Irvine’s Center for Clinical Research.

### MRI Acquisition

Magnetic resonance imaging (MRI) scanning sessions at the University of California, Santa Barbara and Irvine were conducted on 3T Prisma scanners equipped with 64-channel phased-array head/neck coil (of which 50 coils are used for axial brain imaging). High-resolution anatomical scans were acquired using a T1-weighted (T1w) magnetization prepared rapid gradient echo (MPRAGE) sequence (TR = 2500 ms, TE = 2.31 ms, T1 = 934 ms, flip angle = 7°, 0.8 mm thickness) followed by a gradient echo fieldmap (TR = 758 ms; TE1 = 4.92 ms; TE2 = 7.38 ms; flip angle = 60°). A T2-weighted (T2w) turbo spin echo (TSE) scan was also acquired with an oblique coronal orientation positioned orthogonally to the main axis of the hippocampus (TR/TE = 9860/50 ms, flip angle = 122°, 0.4 × 0.4 mm^2^ in-plane resolution, 2 mm slice thickness, 38 interleaved slices with no gap, total acquisition time = 5:42 min). The DSI protocol sampled the entire brain with the following parameters: single phase, TR = 4300 ms, echo time = 100.2 ms, 139 directions, b-max = 4990, FoV = 259 × 259 mm, 78 slices, 1.7986 × 1.7986 × 1.8 mm voxel resolution. These images were linearly registered to the whole-brain T1w MPRAGE image. A custom foam headcase was used to provide extra padding around the head and neck, as well as to minimize head motion. Additionally, a custom-built sound-absorbing foam girdle was placed around the participant’s waist to attenuate sound near the fetus during second and third trimester scanning.

### Image Processing

#### Cortical Volume and Thickness

Cortical thickness and gray matter volume were measured with Advanced Normalization Tools version 2.1.0 (ANTs) (Avants et al., 2011). We first built a subject-specific template (SST) (*antsMultivariateTemplateConstruction2*) and tissue priors (*antsCookTemplatePriors*) based on our subject’s two pre-conception whole-brain T1-weighted scans to examine neuroanatomical changes relative to the subject’s pre-pregnancy baseline. We used labels from the OASIS population template, provided by ANTs, as priors for this step. For each session, the structural image was processed and registered to the SST using the ANTs cortical thickness pipeline (*antsCorticalThickness*). This begins with an N4 bias field correction for field inhomogeneity, then brain extraction using a hybrid registration/segmentation method (see Tustison et al., 2014). Tissue segmentation was performed using Atropos (Avants et al., 2011) to create tissue masks of cerebrospinal fluid, gray matter, white matter, and deep gray matter. Atropos allows prior knowledge to guide the segmentation algorithm, and we used labels from our SST as priors to minimize warping and remain in native subject space. Cortical thickness measurements were then estimated using the DiReCT algorithm (Das et al., 2009), which estimates the gray–white matter interface and the gray matter–CSF interface and computes a diffeomorphic mapping between the two interactions, from which thickness is derived. Each gray matter tissue mask was normalized to the template and multiplied to a Jacobian image that was computed via affine and non-linear transforms. Summary, regional-level estimates of CT, GMV, and CSF for each scan were obtained by taking the first eigenvariate (akin to a ‘weighted mean’, Friston et al., 2006) across all voxels within each parcel of the Schaefer 400-region atlas (Schaefer et al., 2018). We then averaged ROIs across networks, which were defined by the 17-network Schaefer scheme (Yeo et al., 2011; Schaefer et al., 2018). Global measures of CT, GMV, and CSF were computed for each session by summing across all voxels within the respective output image; total brain volume was computed by summing across all voxels within each session’s brain extraction mask. Our findings held when using an SST derived from all 26 MRIs (pre-through postpartum), as well as when estimating the mean (vs. weighted mean) of all voxels within each parcel. The ANTs CT pipeline is highly validated with good test-retest reproducibility and improved ability to predict variables such as age and gender from region-wise CT measurements compared to surface-based FreeSurfer (Tustison et al., 2014). However, to reproduce our findings across software packages, we also ran the T1w data through the longitudinal FreeSurfer cortical thickness pipeline (Dale et al., 1999; Reuter et al., 2012), which corroborated our findings using both the Schaefer-400 (**Fig. S2**; **Tables S1**, **S4**) and popular Desikan-Killiany (Desikan et al., 2004; **Table S3**) cortical parcellations. Whole-brain T1w-based subcortical volume estimates (including cerebellum and lateral ventricles) were also derived using this FreeSurfer pipeline, wherein we derived 28 region-of-interest estimates via the commonly used ‘aseg’ parcellation scheme (Fischl et al., 2002; **Fig. S6A**). A complete reporting of findings can be found in **Supplementary File 2**.

Mean framewise displacement (FWD) estimates from gestation sessions with a 10-minute resting state scan (n = 17) were used to *indirectly* assess whether motion increased throughout pregnancy. Average FWD (millimeters) was extremely minimal across the entire experiment (*M* = 0.13, *SD* = 0.02, *range* = 0.09–0.17) and varied only slightly by pregnancy stage (pre: *M* = 0.11, *SD* = 0.004; first: *M* = 0.11, *SD* = 0.01; second: *M* = 0.14, *SD* = 0.02; third: *M* = 0.16, *SD* = 0.007; post: *M* = 0.13, *SD* = 0.01). While mean FWD did correspond with gestation week (*r* = 0.90, *p* < .001), controlling for this did not alter our main findings (e.g., total GMV negatively associated with gestation; partial correlation: *r* = -0.64, *p* = 0.004) owing to the fact that motion differences between stages were minuscule (**Fig. S4A**).

As a further test of the robustness of the dataset, we ran quality control (QC) assessments on all T1w images using the IQMs pipeline from *MRIQC* (Esteban et al., 2017a). Assessments of interest included 1) coefficient of joint variation (CJV), 2) signal-to-noise ratio for gray matter (SNR), and 3) contrast-to-noise ratios (CNR). All QC metrics fell within expected standard ranges (**Fig. S4B–D**) (Esteban et al., 2017b). Although relationships existed between gestation week and QC measures (CJV, *r* = 0.70, *p* < .001; SNR and CNR, *r* = -0.83, *p* < .001), including these variables in the regression models did not detract from our finding suggesting cortical GMV reductions occur over gestation, especially within regions belonging to attention and somatosensory networks (see **Fig. S5**). When looking across all *MRIQC* outputs, discrepancies were noted in session seven (gestation week nine, first trimester). Removing this day from the analyses only strengthened observed relationships between cortical volume and gestation; however for completeness, data from this day is included in the main findings. These quality control outputs for each session of the experiment can be found in **Supplementary File 2**. Finally, we used FreeSurfer’s Eueler number to evaluate a field-standard *quantitative* assessment of each T1w structural image (Rosen et al., 2018). We observed no significant relationships between the Euler number and gestation week or summary brain metrics. A discrepancy (e.g., 2 SD below average) was noted in session eight; however, removing this session did not detract from our main findings showing reductions in gray matter volume over gestation.

#### Hippocampal Segmentation

T1- and T2-weighted images (n = 25) were submitted to the automatic segmentation of hippocampal subfields package (ASHS) (Yushkevich et al., 2015) for parcellation of seven MTL subregions: CA1, CA2/3, dentate gyrus (DG), subiculum (SUB), perirhinal cortex (PRC), entorhinal cortex (ERC), and parahippocampal cortex (PHC) (see **Fig. S6B**). The ASHS segmentation pipeline automatically segmented the hippocampus in the T2w MRI scans using a segmented population atlas, the Princeton Young Adult 3T ASHS Atlas template (n = 24, mean age 22.5 years; Aly and Turk-Browne, 2016). A rigid-body transformation aligned each T2w image to the respective T1w scan for each day. Using ANTs deformable registration, the T1w was registered to the population atlas. The resulting deformation fields were used to resample the data into the space of the left and right template MTL regions of interest (ROI). Within each template ROI, each of the T2w scans of the atlas package was registered to that day’s T2w scan. The manual atlas segmentations were then mapped into the space of the T2w scan, with segmentation of the T2w scan computed using joint label fusion (Wang et al., 2012). Finally, the corrective learning classifiers contained in ASHS were applied to the consensus segmentation produced by joint label fusion. The output of this step is a corrected segmentation of the T2w scan. Further description of the ASHS protocol can be found in (Yushkevich et al., 2015). T2w scans and segmentations were first visually examined using ITK-SNAP (Yushkevich et al., 2006) for quality assurance and then subjected to manual editing in native space using ITK-SNAP (v.3.8.0-b; author C.M.T.). One session (scan 15, third trimester) was discarded due to erroneous scan orientation. The anterior extent of the segmented labels was anchored 4 mm (2 slices) anterior to the appearance of the limen insulae, and the posterior extent was anchored to the disappearance of hippocampal gray matter from the trigone of the lateral ventricle. Boundaries between perirhinal, entorhinal, and parahippocampal cortices were established in keeping with the Olsen-Amaral-Palombo (OAP) segmentation protocol (Palombo et al., 2013). In instances where automatic segmentation did not clearly correspond to the underlying neuroanatomy, such as when a certain label was missing several gray matter voxels, manual retouching allowed for individual voxels to be added or removed. All results are reported using the manually retouched subregion volumes to ensure the most faithful representation of the underlying neuroanatomy. Scans were randomized and segmentation was performed in a random order, blind to pregnancy stage. To assess intra-rater reliability for the present analyses, two days underwent manual editing a second time. The generalized Dice similarity coefficient (Crum et al., 2006) across subregions was 0.87 and the Intraclass Correlation Coefficient was 0.97, suggesting robust reliability in segmentation.

#### White Matter Microstructure

Diffusion scans were preprocessed using the automation software QSIprep version 0.15.3 (Cieslak et al., 2022) and run primarily with the default parameters, with the exceptions ‘–output resolution 1.8’, ‘–dwi denoise window 5’,–force-spatial-normalization’, ‘–hmc model 3dSHORE’, ‘–hmc-transform Rigid’, and ‘–shoreline iters 2’. Twenty-one sessions were preprocessed and analyzed, with the remaining five scans excluded due to missing data or the corresponding field map for distortion correction. T1w images were corrected for intensity non-uniformity (*N4BiasFieldCorrection*) and skull-stripped (*antsBrainExtraction*). The images underwent spatial normalization and registration to the ICBM 152 Nonlinear Asymmetrical Template. Finally, brain tissue segmentation of CSF, GM, and WM was performed on each brain-extracted T1w using FMRIB’s Automated Segmentation Tool (FAST). Preprocessing of diffusion images began by implementing MP-PCA denoising with a 5-voxel window using MRtrix3’s *dwidenoise* function. B1 field inhomogeneity was corrected using *dwibiascorrect* from MRtrix3 with the N4 algorithm. Motion was corrected using the *SHORELine* method. Susceptibility distortion correction was based on GRE field maps. Preprocessed Nifti scans were prepared for tractography using DSI Studio version Chen-2022-07-31 (Yeh et al., 2016). Diffusion images were converted to Source Code files using the DSI studio command line *‘--action=src’* and a custom script to convert all images. The diffusion data were reconstructed in MNI space using q-space diffeomorphic reconstruction (Yeh et al., 2011) with a diffusion sampling of 1.25 and output resolution of 1.8mm isotropic. The following output metrics were specified to be included in the output FIB file: QA and mean diffusivity (MD). The quality and integrity of reconstructed images were assessed using *‘QC1: SRC Files Quality Control’*. First, consistency of image dimension, resolution, DWI count, shell count was checked for each image. Second, each image was assessed for the “neighboring DWI Correlation” which calculates the correlation coefficient of low-b DWI volumes that have similar gradient direction. Lower correlation values may indicate issues with the diffusion signal due to artifacts or head motion. Finally, DSI studio performed an outlier check, labelling images as a “low quality outlier” if the correlation coefficient was > 3 standard deviations from the absolute mean. None of our scans were flagged as outliers. The reconstructed subject files were aggregated into one connectometry database per metric.

#### Day2Day control dataset

To compare our findings against a control group of non-pregnant densely-sampled individuals, we utilized the Day2Day dataset (Filevich et al., 2017) which offered comparable whole-brain T1 and T2 MTL scans for eight subjects (2 male) scanned 12–50 times over a period of 2–7 months. Each subject was run through the ANTs CT and ASHS processing pipelines as outlined above (see *Cortical Volume and Thickness and Hippocampal Segmentation*). To note, for each subject we created an SST based on their first two sessions for consistency with the primary dataset; subfield volumes for the T2 MTL scans did not undergo manual retouching. Due to missing header information on the publicly available diffusion scans, we were unable to benchmark our white matter changes with the Day2Day dataset.

## Statistical Analysis

Statistical analyses were conducted in R (sMRI; version 3.4.4) and DSI Studio (dMRI; Chen-2022-07-31).

### Summary Brain Metrics

To reflect the existing literature, we first explored brain metrics across the entire study duration (pre-conception through postpartum, N = 26 scans). When including all sessions, total brain volume, GMV, CT, global QA, ventricle volume and CSF displayed non-linear trends over time; therefore, we used generalized additive models (GAM; cubic spline basis, k = 10, smoothing = GCV), a method of non-parametric regression analysis (R package: *mgcv*), to explore the relationship between summary brain metrics (outcome variables) and gestation week (smooth term). Each model underwent examination (*gam.check* function) to ensure it was correctly specified with regards to 1) the choice of basis dimension (*k*) and 2) the distribution of the model residuals (see *mgcv* documentation; Wood, 2017). The general pattern of results held after toggling model parameters; however, we note the risk of overinterpreting complex models with small sample sizes (see Sullivan et al., 2015). To address overfitting and cross-validate our basis type selection, we also fit the data using nonpenalized general linear models (GLM) with both linear and polynomial terms for gestation week. We compared the performance of each GLM (i.e., models using only a linear term vs. models with polynomial terms) via the Akaike information criterion (AIC), which revealed that cubic models consistently outperformed both linear and quadratic models (AIC_diff_ > 3), providing additional evidence for non-linear changes in structural brain variables over time. Determining whether these patterns replicate in larger cohorts and whether complex models are better suited to capture data patterns across individuals will be a necessary next step.

### Cortical Gray Matter Volume & Cortical Thickness

We then narrowed our analyses to the first 19 sessions (baseline – 36 weeks gestation) to assess novel brain changes occurring over gestational window. We first computed Pearson’s product-moment correlation matrices between the following variables: gestation week, estradiol, progesterone, and the 17 network-level average GMV values. We then ran a multivariate regression analysis predicting ROI-level GMV changes by gestation week. To identify which regions were changing at a rate different from the global decrease, we then ran the analyses again to include total GMV in the regression model (globally-corrected data provided in supplementary tables). This was extended to the network level, where we ran partial correlations accounting for total GMV. These same analyses were then run with cortical thickness measures. Percent change at the network level was computed by subtracting the final pregnancy value (36 weeks pregnant) from the first pre-pregnancy baseline, then dividing that difference by said first pre-pregnancy baseline value. All analyses underwent multiple comparisons testing (FDR-corrected at *q* < 0.05).

### Subcortical Gray Matter Volume

A similar statistical approach was taken for subcortical volume estimates. We ran a multivariate regression analysis predicting GMV changes over gestation in 28 regions-of-interest (Fig. S6A) by gestation week (FDR-corrected at q < 0.05).

To evaluate the relationship between gestation week and medial temporal lobe (MTL) subregion volume over pregnancy (n = 7 bilateral subregions; n = 18 MTL scans), we used a combination of linear and non-linear models based on individual subregion data patterns. Models were compared for best fit with each subregion via AIC from the GLM output (as described above). A linear regression model was most appropriate for PHC (AIC_diff_ < 3), whereas a quadratic model performed best for CA1 and CA2/3. As a control, we repeated the analyses with MTL subregion volumes after proportional volume correction of total gray matter volume calculated by ASHS. Finally, we evaluated the relationship between endogenous sex hormones (estrogen and progesterone) and subregion volumes using linear regression. Relationships were considered significant only if they met FDR correction at *q* < .05.

### Diffusion - White Matter Microstructure

DSI Studio’s correlational tractography (Yeh et al., 2016) was used to analyze the relationship between white matter structure and gestational week (n = 15). A truncated model was run to examine the relationship between white matter and sex steroid hormones (n = 10) for the subset of diffusion scans with paired endocrine data. A non-parametric Spearman correlation was used to derive the correlation between gestational week and endocrine factors and our metrics of interest (QA and MD; see **Table S9** and **Fig. S10** for MD results) because the data were not normally distributed. Statistical inference was reached using connectometry, a permutation-based approach that tests the strength of coherent associations found between the local connectome and our variables of interest. It provides higher reliability and replicability by correcting for multiple comparisons. This technique provides a high-resolution characterization of local axonal orientation. The correlational tractography was run with the following parameters: T-score threshold of 2.5, 4 pruning iterations, and a length threshold of 25 voxel distance. To estimate the false discovery rate (FDR), a total of 4000 randomized permutations were applied to obtain the null distribution of the track length. Reported regions were selected based on FDR cutoff (FDR < 0.2, suggested by DSI Studio), and contained at least 10 tracts. For visualization of global QA at each gestational stage, QA values were extracted using DSI Studio’s whole brain fiber tracking algorithm and tractometry (Yeh et al., 2016).

### Day2Day dataset – Measurement Variability

In order to establish a marker of normative variability over the course of half a year, we computed metrics of measurement variability using the Day2Day dataset (Filevich et al., 2017), which provided both whole-brain T1 and high-resolution T2 MTL scans. For each region, *j*, of the Schaefer parcellation, we assessed across-session variability, ε, as:

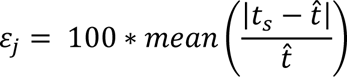

Where *t_s_* is the morphometric measurement of a parcel for session s, and 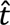 is the mean of *t* across sessions (Jovicich et al., 2013; Tustison et al., 2014). Thus, we defined variability as the mean absolute percent difference between each individual and the mean across sessions. Across-session variability estimates for all 400 regions were then averaged across 8 subjects, and a global measure of cortical GMV variability was computed by averaging across the 400 regions. This approach was repeated independently for the T2 hippocampal scans, wherein we computed across-session variability for each parcel of the ASHS parcellation scheme (n = 7 bilateral subfields). However, it is important to note that *raw* subfield values (i.e., no manual retouching) were used for Day2Day variability assessments and should be interpreted with caution. Finally, to better compare against our own data, we repeated this approach using our participant’s first two baseline scans (i.e., pre-conception) to derive within-subject variability estimates.

Benchmarking our data in this way allows us to capture the degree of change expected due to factors such as image processing and instrumentation variability or other day-to-day changes that could potentially modulate brain size and shape (see Hedges et al., 2022 for review). The percent change observed over pregnancy (baseline vs. 36 weeks gestation) far exceed the expected variability estimated using both the Day2Day dataset (Fig. S11) and our within-subject control data. This was quantified by dividing the observed percent change in GMV metrics (baseline vs. 36 weeks) by the global measure of GMV percent variability of each control group (i.e., Day2Day, within-subject control), independently for cortex and subcortex.

## Acknowledgements

We would like to thank Mario Mendoza for his phlebotomy and MRI assistance at the UCSB Brain Imaging Center. We would like to thank Craig Stark and Rongwen Tain for MRI assistance at the UCI Facility for Imaging and Brain Research (FIBRE), and UCI Institute for Clinical and Translational Science for phlebotomy assistance. We would like to thank Bailey Tranquada-Torres, Rob Woodry, and Nikki Hatamian for their assistance with data collection, as well as Pantea Mozayeni, Candice Taylor, B. Peng, and P.J.C. Peng for their support. Thank you to Dr. Simone Kühn and colleagues for creating the Day2Day dataset and sharing it with our team. Lastly, we would like to thank Magdalena Martínez-García, Susana Carmona, Scott Grafton, Javier Gonzalez-Castillo and Peter Bandettini for their insightful discussions and feedback on this project.

## Data Availability Statement

The dataset consists of 26 MRI scans (T1w, T2w, and diffusion scans) alongside state-dependent measures and serum assessments of ovarian sex hormones for each session. The data will be publicly available on https://openneuro.org/ upon publication. No custom code was used.

## References

1. Ammari, R., Monaca, F., Cao, M., Nassar, E., Wai, P., Del Grosso, N. A., … & Kohl, J. (2023). Hormone-mediated neural remodeling orchestrates parenting onset during pregnancy. Science, 382(6666), 76–81.

2. Barba-Müller, E., Craddock, S., Carmona, S., & Hoekzema, E. (2019). Brain plasticity in pregnancy and the postpartum period: links to maternal caregiving and mental health. Archives of women’s mental health, 22, 289–299.

3. Barrière, D. A., Ella, A., Szeremeta, F., Adriaensen, H., Même, W., Chaillou, E., Migaud, M., Même, S., Lévy, F., & Keller, M. (2021). Brain orchestration of pregnancy and maternal behavior in mice: A longitudinal morphometric study. NeuroImage, 230, 117776.

4. Barth, C., & de Lange, A.-M. G. (2020). Towards an understanding of women’s brain aging: The immunology of pregnancy and menopause. Frontiers in Neuroendocrinology, 58, 100850.

5. Been, L. E., Sheppard, P. A., Galea, L. A., & Glasper, E. R. (2022). Hormones and neuroplasticity: A lifetime of adaptive responses. Neuroscience & Biobehavioral Reviews, 132, 679–690

6. Bethlehem, R. A., Seidlitz, J., White, S. R., Vogel, J. W., Anderson, K. M., Adamson, C., … & Schaare, H. L. (2022). Brain charts for the human lifespan. Nature, 604(7906), 525–533.

7. Blakemore, S. J., & Choudhury, S. (2006). Development of the adolescent brain: implications for executive function and social cognition. Journal of child psychology and psychiatry, 47(3-4), 296–312.

8. Blakemore, S. J., Burnett, S., & Dahl, R. E. (2010). The role of puberty in the developing adolescent brain. Human brain mapping, 31(6), 926–933.

9. Brunton, P. J., & Russell, J. A. (2008). The expectant brain: Adapting for motherhood. Nature Reviews Neuroscience, 9(1), 11–25.

10. Carmona, S., Martínez-García, M., Paternina-Die, M., Barba-Müller, E., Wierenga, L. M., Alemán Gómez, Y., Pretus, C., Marcos-Vidal, L., Beumala, L., Cortizo, R., Pozzobon, C., Picado, M., Lucco, F., García-García, D., Soliva, J. C., Tobeña, A., Peper, J. S., Crone, E. A., Ballesteros, A., … Hoekzema, E. (2019). Pregnancy and adolescence entail similar neuroanatomical adaptations: A comparative analysis of cerebral morphometric changes. Human Brain Mapping, 40(7), 2143–2152.

11. Celik, A., Somer, M., Kukreja, B., Wu, T., & Kalish, B. T. (2022). The Genomic Architecture of Pregnancy-Associated Plasticity in the Maternal Mouse Hippocampus. Eneuro, 9(5).

12. Chaker, Z., Segalada, C., Kretz, J. A., Acar, I. E., Delgado, A. C., Crotet, V., … & Doetsch, F. (2023). Pregnancy-responsive pools of adult neural stem cells for transient neurogenesis in mothers. Science, 382(6673), 958–963.

13. Crespi, C., Cerami, C., Dodich, A., Canessa, N., Arpone, M., Iannaccone, S., … & Cappa, S. F. (2014). Microstructural white matter correlates of emotion recognition impairment in Amyotrophic Lateral Sclerosis. cortex, 53, 1–8.

14. de Lange, A.-M. G., Kaufmann, T., van der Meer, D., Maglanoc, L. A., Alnæs, D., Moberget, T., Douaud, G., Andreassen, O. A., & Westlye, L. T. (2019). Population-based neuroimaging reveals traces of childbirth in the maternal brain. Proceedings of the National Academy of Sciences, 116(44), 22341–22346.

15. Deligiannidis, K. M., Meltzer-Brody, S., Maximos, B., Peeper, E. Q., Freeman, M., Lasser, R., … & Doherty, J. (2023). Zuranolone for the Treatment of Postpartum Depression. American Journal of Psychiatry, appi-ajp.

16. Diamond, M. C., Johnson, R. E., & Ingham, C. (1971). Brain plasticity induced by environment and pregnancy. International Journal of Neuroscience, 2(4-5), 171–178.

17. Duarte-Guterman, P., Richard, J. E., Lieblich, S. E., Eid, R. S., Lamers, Y., & Galea, L. A. M. (2023). Cellular and molecular signatures of motherhood in the adult and ageing rat brain. Open Biology, 13(11), 230217.

18. Dulac, C., O’Connell, L. A., & Wu, Z. (2014). Neural control of maternal and paternal behaviors. Science, 345(6198), 765–770.

19. Filevich, E., Lisofsky, N., Becker, M., Butler, O., Lochstet, M., Martensson, J., … & Kühn, S. (2017). Day2day: investigating daily variability of magnetic resonance imaging measures over half a year. BMC neuroscience, 18, 1–8.

20. Haim, A., Julian, D., Albin-Brooks, C., Brothers, H. M., Lenz, K. M., & Leuner, B. (2017). A survey of neuroimmune changes in pregnant and postpartum female rats. Brain, behavior, and immunity, 59, 67–78.

21. Herbet, G., Zemmoura, I., & Duffau, H. (2018). Functional anatomy of the inferior longitudinal fasciculus: from historical reports to current hypotheses. Frontiers in neuroanatomy, 12, 77.

22. Hoekzema, E., Barba-Müller, E., Pozzobon, C., Picado, M., Lucco, F., García-García, D., Soliva, J. C., Tobeña, A., Desco, M., Crone, E. A., Ballesteros, A., Carmona, S., & Vilarroya, O. (2017). Pregnancy leads to long-lasting changes in human brain structure. Nature Neuroscience, 20(2), 287–296.

23. Hoekzema, E., van Steenbergen, H., Straathof, M., Beekmans, A., Freund, I. M., Pouwels, P. J. W., & Crone, E. A. (2022). Mapping the effects of pregnancy on resting state brain activity, white matter microstructure, neural metabolite concentrations and grey matter architecture. Nature Communications, 13(1), 6931.

24. Kohl, J., Babayan, B. M., Rubinstein, N. D., Autry, A. E., Marin-Rodriguez, B., Kapoor, V., Miyamishi, K., Zweifel, L. S., Luo, L., Uchida, N., & Dulac, C. (2018). Functional circuit architecture underlying parental behaviour. Nature, 556(7701), Article 7701.

25. Lövdén, M., Bodammer, N. C., Kühn, S., Kaufmann, J., Schütze, H., Tempelmann, C., … & Lindenberger, U. (2010). Experience-dependent plasticity of white-matter microstructure extends into old age. Neuropsychologia, 48(13), 3878–3883.

26. Makris, N., Oscar-Berman, M., Jaffin, S. K., Hodge, S. M., Kennedy, D. N., Caviness, V. S., … & Harris, G. J. (2008). Decreased volume of the brain reward system in alcoholism. Biological psychiatry, 64(3), 192–202.

27. Martínez-García, M., Paternina-Die, M., Desco, M., Vilarroya, O., & Carmona, S. (2021a). Characterizing the Brain Structural Adaptations Across the Motherhood Transition. Frontiers in Global Women’s Health, 2(742775).

28. Martínez-García, M., Paternina-Die, M., Barba-Müller, E., Martín de Blas, D., Beumala, L., Cortizo, R., Pozzobon, C., Marcos-Vidal, L., Fernández-Pena, A., & Picado, M. (2021b). Do Pregnancy-Induced Brain Changes Reverse? The Brain of a Mother Six Years after Parturition. Brain Sciences, 11(2), 168.

29. Martínez-García, M., Jacobs E.G., de Lange Ann Marie, Carmon S. Advancing the neuroscience of human pregnancy. In press.

30. Orchard, E. R., Ward, P. G. D., Chopra, S., Storey, E., Egan, G. F., & Jamadar, S. D. (2020). Neuroprotective Effects of Motherhood on Brain Function in Late Life: A Resting-State fMRI Study. Cerebral Cortex, 1270–1283.

31. Orchard, E. R., Rutherford, H. J. V., Holmes, A. J., & Jamadar, S. D. (2023). Matrescence: Lifetime impact of motherhood on cognition and the brain. Trends in Cognitive Sciences.

32. Paternina-Die, M., Martínez-García, M., Martín de Blas, D., Noguero, I., Servin-Barthet, C., Pretus, C., … & Carmona, S. (2024). Women’s neuroplasticity during gestation, childbirth and postpartum. Nature Neuroscience, 1–9.

33. Pawluski, Jodi L., Joseph S. Lonstein, and Alison S. Fleming. “The neurobiology of postpartum anxiety and depression.” Trends in Neurosciences 40.2 (2017): 106–120.

34. Pawluski, J. L., Hoekzema, E., Leuner, B., & Lonstein, J. S. (2022). Less Can Be More: Fine Tuning the Maternal Brain. Neuroscience & Biobehavioral Reviews.

35. Pritschet, L., Taylor, C. M., Santander, T., & Jacobs, E. G. (2021). Applying dense-sampling methods to reveal dynamic endocrine modulation of the nervous system. Current opinion in behavioral sciences, 40, 72–78.

36. Puri, T. A., Richard, J. E., & Galea, L. A. M. (2023). Beyond sex differences: Short- and long-term effects of pregnancy on the brain. Trends in Neurosciences.

37. Rodrigo, M. J., León, I., Góngora, D., Hernández-Cabrera, J. A., Byrne, S., & Bobes, M. A. (2016). Inferior fronto-temporo-occipital connectivity: a missing link between maltreated girls and neglectful mothers. Social cognitive and affective neuroscience, 11(10), 1658–1665.

38. Servin-Barthet, C., Martínez-García, M., Pretus, C., Paternina-Die, M., Soler, A., Khymenets, O., … & Carmona, S. (2023). The transition to motherhood: linking hormones, brain and behaviour. Nature Reviews Neuroscience, 24(10), 605–619

39. Shehata, H. A., & Okosun, H. (2004). Neurological disorders in pregnancy. Current Opinion in Obstetrics and Gynecology, 16(2), 117–122.

40. Spalek, K., Straathof, M., Koyuncu, L., Grydeland, H., van der Geest, A., van‘t Hof, S. R., … & Hoekzema, E. (2024). Pregnancy renders anatomical changes in hypothalamic substructures of the human brain that relate to aspects of maternal behavior. Psychoneuroendocrinology, 107021.

41. Taxier, L. R., Gross, K. S., & Frick, K. M. (2020). Oestradiol as a neuromodulator of learning and memory. Nature Reviews Neuroscience, 21(10), 535–550.

42. Thornburg, K. L., Bagby, S. P., & Giraud, G. D. (2015). Maternal Adaptations to Pregnancy. In Knobil and Neill’s Physiology of Reproduction (pp. 1927–1955). Elsevier, Inc.

43. Tooley, U.A., Bassett, D.S., & Mackey, A.P. (2021). Environmental influences on the pace of brain development. Nature Reviews Neuroscience, 22, 372–384.

44. Wang, Y., Metoki, A., Alm, K. H., & Olson, I. R. (2018). White matter pathways and social cognition. Neuroscience & Biobehavioral Reviews, 90, 350–370.

45. Wang, Z., Liu, J., Shuai, H., Cai, Z., Fu, X., Liu, Y., … & Yang, B. X. (2021). Mapping global prevalence of depression among postpartum women. Translational psychiatry, 11(1), 543.

46. World Health Organization (2022). “Maternal, newborn, child and adolescent health and ageing*”*

47. Zekelman, L. R., Zhang, F., Makris, N., He, J., Chen, Y., Xue, T., … & O’Donnell, L. J. (2022). White matter association tracts underlying language and theory of mind: An investigation of 809 brains from the Human Connectome Project. Neuroimage, 246, 118739.

## References

1. Aly, M., & Turk-Browne, N. B. (2016). Attention stabilizes representations in the human hippocampus. Cerebral Cortex, 26(2), 783–796.

2. Avants, B. B., Tustison, N. J., Song, G., Cook, P. A., Klein, A., & Gee, J. C. (2011). A reproducible evaluation of ANTs similarity metric performance in brain image registration. Neuroimage, 54(3), 2033–2044.

3. Buysse, D. J., Reynolds III, C. F., Monk, T. H., Berman, S. R., & Kupfer, D. J. (1989). The Pittsburgh Sleep Quality Index: a new instrument for psychiatric practice and research. Psychiatry research, 28(2), 193–213.

4. Cieslak, M., Cook, P. A., He, X., Yeh, F. C., Dhollander, T., Adebimpe, A., … & Satterthwaite, T. D. (2021). QSIPrep: an integrative platform for preprocessing and reconstructing diffusion MRI data. Nature methods, 18(7), 775–778.

5. Cohen, S., Kamarck, T., & Mermelstein, R. (1983). A global measure of perceived stress. Journal of health and social behavior, 385–396.

6. Crum, W. R., Camara, O., & Hill, D. L. (2006). Generalized overlap measures for evaluation and validation in medical image analysis. IEEE transactions on medical imaging, 25(11), 1451–1461.

7. Dale, A. M., Fischl, B., & Sereno, M. I. (1999). Cortical surface-based analysis: I. Segmentation and surface reconstruction. Neuroimage, 9(2), 179–194.

8. Das, S. R., Avants, B. B., Grossman, M., & Gee, J. C. (2009). Registration based cortical thickness measurement. Neuroimage, 45(3), 867–879.

9. Desikan, R. S., Ségonne, F., Fischl, B., Quinn, B. T., Dickerson, B. C., Blacker, D., … & Killiany, R. J. (2006). An automated labeling system for subdividing the human cerebral cortex on MRI scans into gyral based regions of interest. Neuroimage, 31(3), 968–980.

10. Esteban, O., Birman, D., Schaer, M., Koyejo, O. O., Poldrack, R. A., & Gorgolewski, K. J. (2017a). MRIQC: Advancing the automatic prediction of image quality in MRI from unseen sites. PloS one, 12(9), e0184661.

11. Esteban, O., Blair, R.W., Nielson, J.C., Varada, S., Marrett, S., Thomas, A.G., Poldrack, R.A., & Gorgolewski, K.J. (2017b); doi: 10.1101/216671

12. Fischl, B., Salat, D. H., Busa, E., Albert, M., Dieterich, M., Haselgrove, C., … & Dale, A. M. (2002). Whole brain segmentation: automated labeling of neuroanatomical structures in the human brain. Neuron, 33(3), 341–355.

13. Friston, K. J., Rotshtein, P., Geng, J. J., Sterzer, P., & Henson, R. N. (2006). A critique of functional localisers. Neuroimage, 30(4), 1077–1087.

14. Hedges, E. P., Dimitrov, M., Zahid, U., Vega, B. B., Si, S., Dickson, H., … & Kempton, M. J. (2022). Reliability of structural MRI measurements: The effects of scan session, head tilt, inter-scan interval, acquisition sequence, FreeSurfer version and processing stream. Neuroimage, 246, 118751.

15. Jovicich, J., Marizzoni, M., Sala-Llonch, R., Bosch, B., Bartrés-Faz, D., Arnold, J., … & PharmaCog Consortium. (2013). Brain morphometry reproducibility in multi-center 3 T MRI studies: a comparison of cross-sectional and longitudinal segmentations. Neuroimage, 83, 472–484.

16. Mowinckel, A. M., & Vidal-Piñeiro, D. (2020). Visualization of brain statistics with R packages ggseg and ggseg3d. Advances in Methods and Practices in Psychological Science, 3(4), 466–483.

17. Palombo, D. J., Amaral, R. S., Olsen, R. K., Müller, D. J., Todd, R. M., Anderson, A. K., & Levine, B. (2013). KIBRA polymorphism is associated with individual differences in hippocampal subregions: evidence from anatomical segmentation using high-resolution MRI. Journal of Neuroscience, 33(32), 13088–13093.

18. Pollock, V., Cho, D. W., Reker, D., & Volavka, J. (1979). Profile of mood states: the factors and their physiological correlates. The Journal of nervous and mental disease, 167(10), 612–614.

19. Reuter, M., Schmansky, N. J., Rosas, H. D., & Fischl, B. (2012). Within-subject template estimation for unbiased longitudinal image analysis. Neuroimage, 61(4), 1402–1418.

20. Rosen, A. F., Roalf, D. R., Ruparel, K., Blake, J., Seelaus, K., Villa, L. P., … & Satterthwaite, T. D. (2018). Quantitative assessment of structural image quality. Neuroimage, 169, 407–418.

21. Schaefer, A., Kong, R., Gordon, E. M., Laumann, T. O., Zuo, X. N., Holmes, A. J., … & Yeo, B. T. (2018). Local-global parcellation of the human cerebral cortex from intrinsic functional connectivity MRI. Cerebral cortex, 28(9), 3095–3114.

22. Speilberger, C. D., & Vagg, P. R. (1984). Psychometric properties of the STAI: a reply to Ramanaiah, Franzen, and Schill. Journal of personality assessment, 48(1), 95–97.

23. Sullivan, K. J., Shadish, W. R., & Steiner, P. M. (2015). An introduction to modeling longitudinal data with generalized additive models: applications to single-case designs. Psychological methods, 20(1), 26.

24. Tustison, N. J., Cook, P. A., Klein, A., Song, G., Das, S. R., Duda, J. T., … & Avants, B. B. (2014). Large-scale evaluation of ANTs and FreeSurfer cortical thickness measurements. Neuroimage, 99, 166–179.

25. Wang, H., Suh, J. W., Das, S. R., Pluta, J. B., Craige, C., & Yushkevich, P. A. (2012). Multi-atlas segmentation with joint label fusion. IEEE transactions on pattern analysis and machine intelligence, 35(3), 611–623.

26. Wood, S. N. (2017). Generalized additive models: an introduction with R. chapman and hall/CRC.

27. Yeh, F. C., & Tseng, W. Y. I. (2011). NTU-90: a high angular resolution brain atlas constructed by q-space diffeomorphic reconstruction. Neuroimage, 58(1), 91–99.

28. Yeh, F. C., Badre, D., & Verstynen, T. (2016). Connectometry: a statistical approach harnessing the analytical potential of the local connectome. Neuroimage, 125, 162–171

29. Yeo, B. T., Krienen, F. M., Sepulcre, J., Sabuncu, M. R., Lashkari, D., Hollinshead, M., … & Buckner, R. L. (2011). The organization of the human cerebral cortex estimated by intrinsic functional connectivity. Journal of neurophysiology, 106(3), 1125–1165.

30. Yushkevich, P. A., Piven, J., Hazlett, H. C., Smith, R. G., Ho, S., Gee, J. C., & Gerig, G. (2006). User-guided 3D active contour segmentation of anatomical structures: significantly improved efficiency and reliability. Neuroimage, 31(3), 1116–1128.

31. Yushkevich, P. A., Pluta, J. B., Wang, H., Xie, L., Ding, S. L., Gertje, E. C., … & Wolk, D. A. (2015). Automated volumetry and regional thickness analysis of hippocampal subfields and medial temporal cortical structures in mild cognitive impairment. Human brain mapping, 36(1), 258–287.

