## Supplementary File 1 for "Neuroanatomical changes observed over the course of a human pregnancy"

**This file includes:** Supplementary Figures (12) and Tables (10)

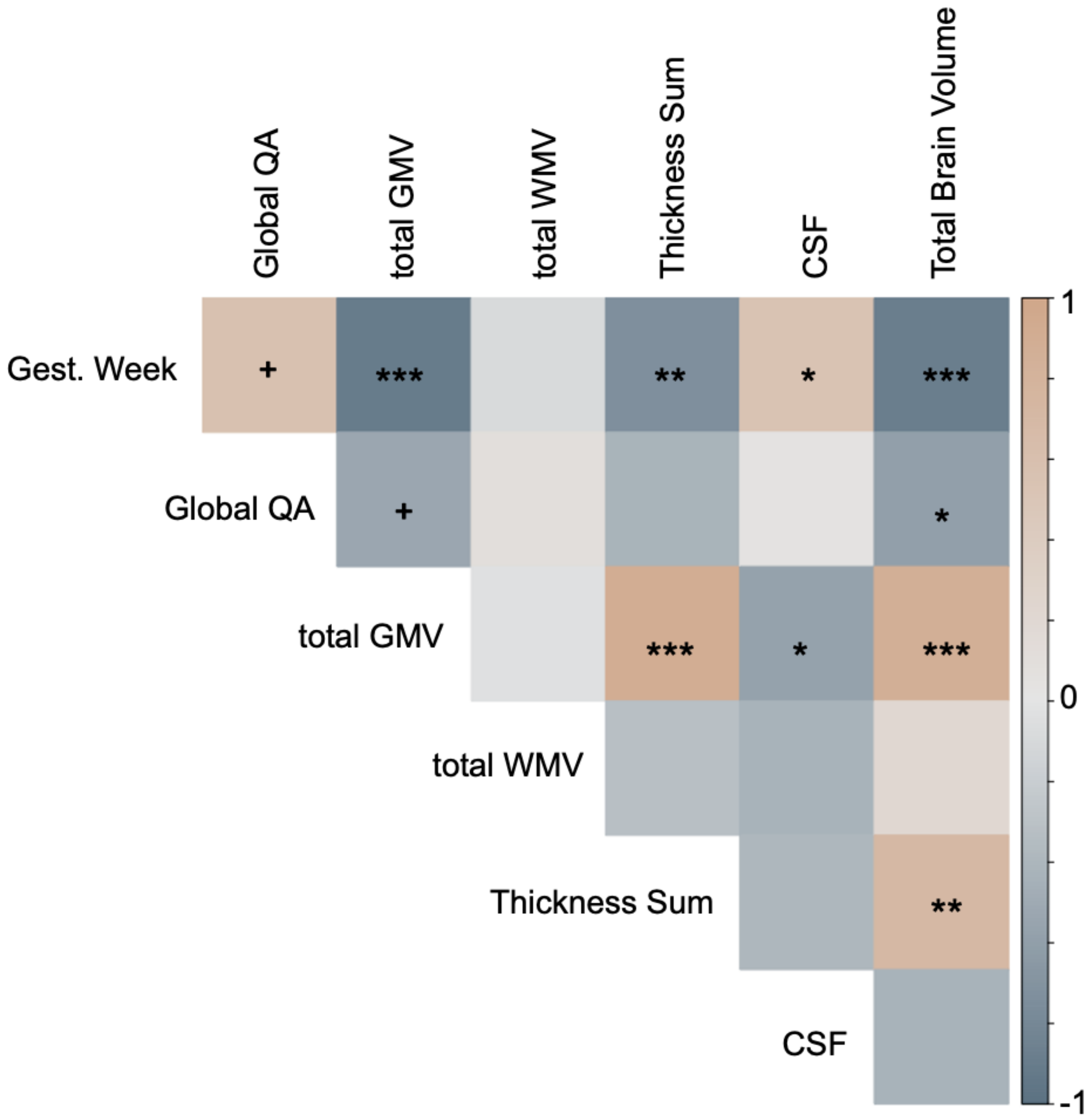


**Supplementary Figure 1.** Relationships between summary (i.e., total) brain measures and gestation week across pregnancy (baseline – 36 weeks). Total brain volume, gray matter volume, and cortical thickness were positively associated with one another, whereas CSF and global QA (weakly) demonstrated negative relationships with gray matter volume. Several summary metrics were not normally distributed—determined via Shapiro-Wilk's test for normality—so Spearman rank correlations were used. *Abbreviations:* QA = global anisotropy; GMV, gray matter volume; WMV, white matter volume; CSF, cerebrospinal fluid. Asterisks indicate significant correlation after FDR-correction ($q < 0.05$). ***$q < 0.001$; **$q < 0.01$, *$q < 0.05$, $^{+}q < 0.10$

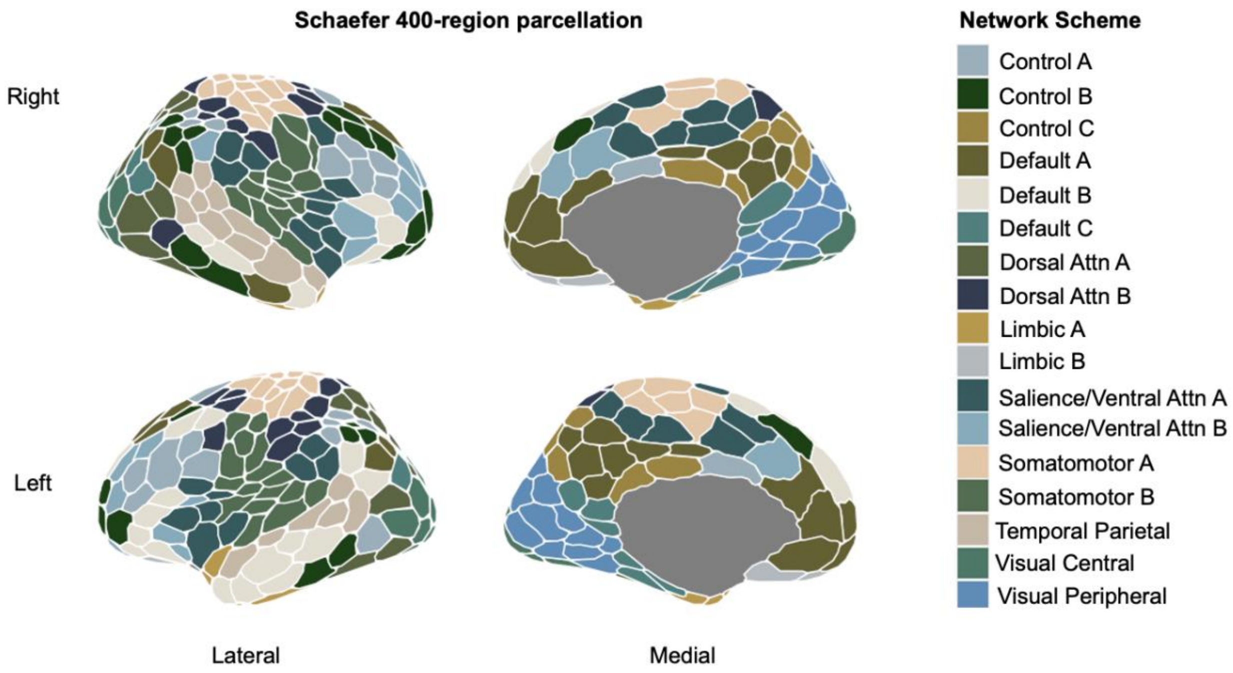


**Supplementary Figure 2.** The brain was parcellated into 400 cortical regions which were then assigned to one of eight networks based on previously identified anatomical and functional associations (Schaefer et al., 2018, Yeo et al., 2011). Colors indicate regional network membership.

**Supplementary Table 1.** Raw gray matter volume and correlations with gestation metrics by network

| Network | Average | Range | Progesterone *N = 13* | 17β-estradiol *N = 15* | Gestation week *N = 19* | Gestation week **controlling for total GMV* | Gestation week *FreeSurfer Results |
|---|---|---|---|---|---|---|---|
| Control A | 0.64 | 0.62 – 0.67 | -0.95*** | -0.88*** | -0.93*** | -0.52* | -0.60** |
| Control B | 0.67 | 0.66 – 0.69 | -0.87*** | -0.88*** | -0.92*** | -0.57* | -0.55* |
| Control C | 0.68 | 0.65 - 0.70 | -0.94*** | -0.90*** | -0.88*** | -0.25 | -0.78*** |
| Default Mode A | 0.70 | 0.69 – 0.72 | -0.95*** | -0.93*** | -0.93*** | -0.57* | -0.76*** |
| Default Mode B | 0.70 | 0.69 – 0.72 | -0.95*** | -0.87*** | -0.91*** | -0.29 | -0.72*** |
| Default Mode C | 0.72 | 0.71 – 0.73 | -0.86*** | -0.72** | -0.69** | 0.77*** | -0.84*** |
| Dorsal Attention A | 0.66 | 0.64 – 0.68 | -0.97*** | -0.89*** | -0.91*** | -0.36 | -0.64** |
| Dorsal Attention B | 0.65 | 0.62 – 0.68 | -0.97*** | -0.91*** | -0.95*** | -0.72*** | -0.55* |
| Salience Ventral Attn A | 0.70 | 0.67 – 0.72 | -0.97*** | -0.91*** | -0.95*** | -0.71*** | -0.92*** |
| Salience Ventral Attn B | 0.72 | 0.70 – 0.73 | -0.92*** | -0.89*** | -0.92*** | -0.41 | -0.69** |
| Somatomotor A | 0.65 | 0.61 – 0.67 | -0.95*** | -0.88*** | -0.93*** | -0.54* | -0.76*** |
| Somatomotor B | 0.70 | 0.67 – 0.73 | -0.95*** | -0.91*** | -0.92*** | -0.52* | -0.91*** |
| Limbic A | 0.76 | 0.74 – 0.79 | -0.11 | 0.16 | 0.14 | 0.65** | -0.08 |
| Limbic B | 0.70 | 0.66 – 0.74 | -0.48 | -0.67** | -0.50* | 0.27 | -0.80*** |
| Temporal Parietal | 0.72 | 0.70 – 0.74 | -0.97*** | -0.92*** | -0.89*** | -0.10 | -0.87*** |
| Visual Central | 0.67 | 0.65 – 0.69 | -0.82*** | -0.78*** | -0.72*** | 0.51* | -0.66** |
| Visual Peripheral | 0.70 | 0.69 – 0.72 | -0.92*** | -0.86*** | -0.70** | 0.71*** | -0.72*** |

FDR-corrected at $q < 0.05$; *$q \leq 0.05$, **$q \leq 0.01$, ***$q \leq 0.001$

**Supplementary Table 1**. Raw gray matter volume (GMV) by network. Pearson's product-moment correlations were used to determine relationships between GMV and gestation week, estradiol, and progesterone (baseline – 36 weeks). The last two columns examine relationships between gestation week and average network GMV after controlling for global shifts in GMV (left) and deriving volumes via FreeSurfer in order to evaluate consistency across software pipelines (right).

**Supplementary Table 2.** Top gray matter volume ROIs associated with gestation week

| Schaefer ROI | Network | Hemisphere | T-stat (raw) | T-stat (partial) |
|---|---|---|---|---|
| PFCd_1 | ContA | Right | -6.36*** | -5.33* |
| IPS_2 | ContA | Left | -4.63*** | -3.47* |
| PFCd_1 | ContB | Left | -4.19*** | -4.74* |
| PFCmp_1 | ContB | Right | -8.38*** | -3.72* |
| PFCmp_1 | ContB | Left | -5.29*** | -3.38* |
| pCun_3 | ContC | Right | -10.10*** | -3.26* |
| IPL_1 | DMNA | Left | -10.94*** | -4.23* |
| PFCd_3 | DMNA | Left | -7.90*** | -4.05* |
| pCunPCC_4 | DMNA | Left | -10.02*** | -3.67* |
| pCunPCC_2 | DMNA | Right | -9.60*** | -3.36* |
| PFCd_2 | DMNA | Right | -8.55*** | -3.33* |
| SPL_3 | DorsAttnA | Left | -10.61*** | -3.59* |
| PostC_6 | DorsAttnB | Right | -13.84*** | -5.35* |
| PostC_3 | DorsAttnB | Left | -14.40*** | -4.95* |
| PostC_7 | DorsAttnB | Right | -7.80*** | -4.56* |
| PostC_7 | DorsAttnB | Left | -9.15*** | -4.48* |
| PostC_9 | DorsAttnB | Left | -8.63*** | -4.08* |
| PostC_8 | DorsAttnB | Left | -8.55*** | -3.96* |
| FEF_2 | DorsAttnB | Left | -8.12*** | -3.77* |
| PostC_8 | DorsAttnB | Right | -9.37*** | -3.75* |
| PostC_1 | DorsAttnB | Left | -12.00*** | -3.53* |
| FEF_3 | DorsAttnB | Left | -10.88*** | -3.41* |
| Ins_1 | SalVentAttnA | Left | -4.90*** | -5.22* |
| FrOper_3 | SalVentAttnA | Right | -11.57*** | -4.26* |
| ParOper_3 | SalVentAttnA | Left | -8.76*** | -3.78* |
| FrMed_2 | SalVentAttnA | Right | -4.21*** | -3.37* |
| Ins_3 | SalVentAttnB | Left | -7.09*** | -5.07* |
| SomMotA_11 | SomMotA | Left | -11.96*** | -4.12* |
| SomMotA_12 | SomMotA | Left | -4.90*** | -4.11* |
| SomMotA_8 | SomMotA | Left | -7.69*** | -3.76* |
| SomMotA_5 | SomMotA | Right | -10.67*** | -3.23* |
| TempPar_3 | TempPar | Left | -11.35*** | -3.40* |
| PHC_2 | DMNC | Right | -2.89* | 4.09* |
| TempPole_3 | LimbicA | Right | 2.78 | 3.54* |
| ExStr_2 | VisCent | Left | -4.71*** | 4.43* |
| ExStrInf_2 | VisPeri | Right | 0.18 | 4.57* |

FDR-corrected at $q < 0.05$; *$q \leq 0.05$, **$q \leq 0.01$, ***$q \leq 0.001$

**Supplementary Table 2**. A subset of regions that demonstrate significant relationships between GMV and gestation week, before (raw) and after controlling for total GMV (partial) in a multivariate regression analysis. Negative ‘partial’ test-statistics suggest these regions-of-interest (ROIs) decrease at a rate greater than the global decrease; positive ‘partial’ test-statistics suggest these ROIs decrease at a rate slower than the global increase. Note, this is not an exhaustive list of results; see **Supplementary File 2** for complete reporting. Abbreviations: PostC = post central; FEF = frontal eye fields; Ins = insula; FrOper = frontal operculum; ParOper = parietal operculum; FrMed = frontal medial; ParOper = parietal operculum; IPL = inferior parietal lobule; PFCd = dorsal prefrontal cortex; pCunPCC = precuneus posterior cingulate cortex; PFCd = dorsal prefrontal cortex; PFCmd = medial posterior prefrontal cortex; IPS = intraparietal sulcus; SPL = superior parietal lobule.

**Supplementary Table 3.** Top Desikan-Killiany GMV ROIs associated with gestation week

| DSK ROI | Hemisphere | T-stat (raw) | T-stat (partial) |
|---|---|---|---|
| Transverse Temporal | Bilateral | L: -8.59***, R: -5.05*** | L: -4.70**, R: -3.80* |
| Lateral Orbital Frontal | Bilateral | L: -5.60***, R: -4.62*** | L: -3.27*, R: -3.51* |
| Medial Orbital Frontal | Right | -6.03*** | -4.80** |
| Lingual | Right | -4.98*** | -3.58* |
| Posterior Cingulate | Left | -7.40*** | -3.55* |
| Pericalcarine | Right | -2.78* | -3.44* |
| Precuneus | Right | -7.76*** | -3.35* |
| Superior Temporal | Left | -7.54*** | -3.26* |
| Insula | Right | -5.14*** | -3.22* |
| Post Central Gyrus | Left | -7.36*** | -3.04* |
| Pars Opercularis | Left | -7.30*** | -3.00* |
| Superior Parietal | Left | -2.80* | 2.94* |
| Inferior Parietal | Right | -0.70 (ns) | 2.97* |
| Rostral Middle Frontal | Bilateral | L: -1.50 (ns), R: -2.48* | L: 3.84*, R: 3.23* |
| Supramarginal | Right | -1.04 (ns) | 4.62** |

FDR-corrected at $q < 0.05$; *$q \leq 0.05$, **$q \leq 0.01$, ***$q \leq 0.001$

**Supplementary Table 3**. Top GMV Desikan-Killiany regions of interest (ROIs) significantly associated with gestation week (raw), including those that remain significant after controlling for total GMV (partial). This allows us to identify regions with GMV decreasing at a rate greater (negative associations) or slower (positive associations) than the global decrease over the gestational window. Volumetric estimates were derived from FreeSurfer to cross-validate with a different pipeline and parcellation. N.b., this is not an exhaustive list; see full results reported in **Supplementary File 2**.

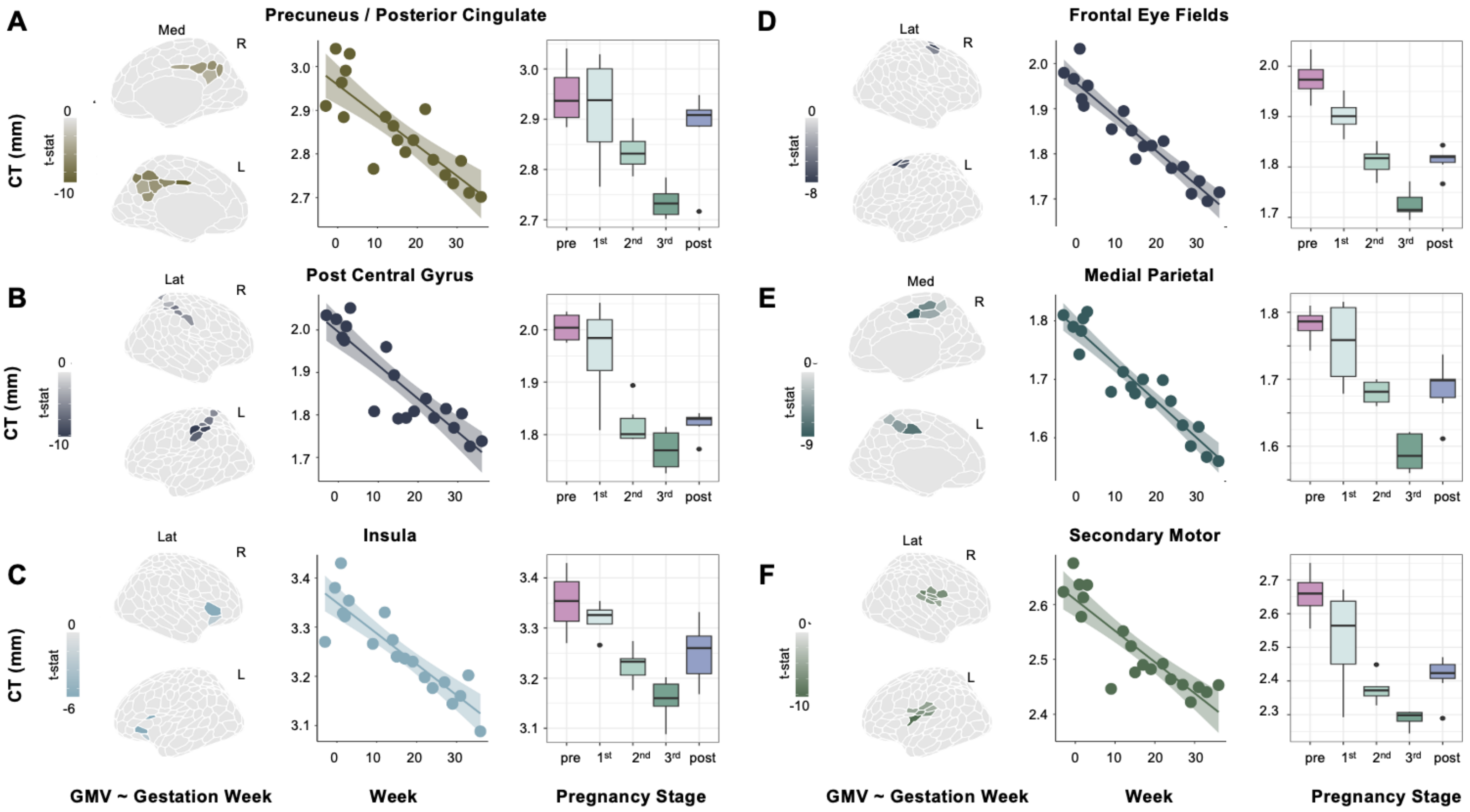


**Supplementary Figure 3.** Six representative regions, color coded by major subnetworks (see **Fig. S2**), that exhibit pronounced cortical thickness change across gestation. **A–F)** For each panel, we display results of a multivariate regression revealing significant associations between regional CT and gestation week (left), Pearson's product-moment correlation between average CT of the regions-of-interest (ROIs) and gestation week (middle; *gestation only*), and summary ROI CT by pregnancy stage across the whole study (right; *gestation and postpartum*). All statistical tests were corrected for multiple comparisons (FDR at $q < 0.05$). N.b., shown here are raw data values (see **Tables S4–5** and **Supplementary File 2** for exhaustive list). Brain visualizations created with R-package *ggseg* (Mowinckel and Vidal-Piñeiro, 2020).

**Supplementary Table 4**. Raw cortical thickness and associations with gestation metrics at network level

| Network | Average | Range | Progesterone *N = 13* | 17β-estradiol *N = 15* | Gestation week *N = 19* | Gestation Week *controlling for total CT* | Gestation Week *FreeSurfer Results* |
|---|---|---|---|---|---|---|---|
| Control A | 1.87 | 1.75 – 2.02 | -0.90*** | -0.80*** | -0.84*** | -0.22 | -0.59* |
| Control B | 2.44 | 2.32 – 2.65 | -0.89*** | -0.85*** | -0.83*** | -0.05 | -0.46 |
| Control C | 2.07 | 1.95 – 2.27 | -0.90*** | -0.80*** | -0.75*** | 0.25 | -0.83*** |
| Default Mode A | 2.69 | 2.57 – 2.91 | -0.91*** | -0.85*** | -0.82*** | -0.01 | -0.83*** |
| Default Mode B | 2.57 | 2.46 – 2.78 | -0.85*** | -0.79*** | -0.80*** | 0.27 | -0.54* |
| Default Mode C | 2.91 | 2.78– 3.05 | -0.49* | -0.49 | -0.40 | 0.86*** | -0.81*** |
| Dorsal Attention A | 2.28 | 2.11 – 2.48 | -0.91*** | -0.82*** | -0.77*** | 0.36 | -0.46 |
| Dorsal Attention B | 1.92 | 1.77 – 2.12 | -0.94*** | -0.88*** | -0.90*** | -0.67* | -0.51 |
| Salience Ventral Attn A | 2.46 | 2.33 – 2.62 | -0.95*** | -0.91*** | -0.91*** | -0.79*** | -0.93*** |
| Salience Ventral Attn B | 2.73 | 2.61 – 2.94 | -0.91*** | -0.86*** | -0.86*** | -0.39 | -0.78** |
| Somatomotor A | 1.76 | 1.52 – 2.23 | -0.86*** | -0.81*** | -0.81*** | -0.16 | -0.62* |
| Somatomotor B | 2.17 | 2.01 – 2.45 | -0.93*** | -0.90*** | -0.89*** | -0.77*** | -0.83*** |
| Limbic A | 3.76 | 3.39 – 3.97 | -0.39 | -0.23 | -0.32 | 0.62* | -0.46 |
| Limbic B | 2.81 | 2.53 – 3.12 | -0.66* | -0.75** | -0.59** | 0.26 | -0.75** |
| Temporal Parietal | 2.66 | 2.46 – 2.86 | -0.94*** | -0.91*** | -0.80*** | -0.03 | -0.73** |
| Visual Central | 2.24 | 1.89 – 2.50 | -0.79** | -0.76** | -0.63** | 0.46 | -0.31 |
| Visual Peripheral | 2.34 | 2.10 – 2.50 | -0.63* | -0.44 | -0.22 | 0.81*** | -0.56* |

FDR-corrected at $q < 0.05$; *$q \leq 0.05$, **$q \leq 0.01$, ***$q \leq 0.001$

**Supplementary Table 4**. Raw cortical thickness (CT) by network. Pearson's product-moment correlations were used to determine relationships between cortical thickness and gestation week, estradiol, and progesterone (baseline – 36 weeks). The last two columns examine relationships between gestation week and average network CT after controlling for global shifts in CT (left) and deriving thickness estimates via FreeSurfer in order to evaluate consistency across software pipelines (right).

**Supplementary Table 5.** Top cortical thickness ROIs associated with gestation week

| Schaefer ROI | Network | Hemisphere | T-stat (raw) | T-stat (partial) |
|---|---|---|---|---|
| PFCl_3 | ContA | Left | -9.90*** | -4.23* |
| PostC_2 | DorsAttnB | Left | -10.20*** | -4.94 |
| PostC_3 | DorsAttnB | Left | -9.50*** | -4.05* |
| FEF_2 | DorsAttnB | Left | -8.86*** | -3.91* |
| FEF_3 | DorsAttnB | Left | -7.80*** | -3.90* |
| FEF_2 | DorsAttnB | Right | -8.63*** | -3.70* |
| ParMed_1 | SalVentAttnA | Right | -11.57*** | -5.72** |
| ParMed_1 | SalVentAttnA | Left | -9.38*** | -4.68* |
| Ins_2 | SalVentAttnB | Left | -6.77*** | -3.77* |
| Ins_1 | SomB | Left | -11.41*** | -5.22* |
| S2_1 | SomB | Left | -10.39*** | -4.72* |
| TempPole_4 | LimbicA | Left | -1.48 | 4.73* |
| ExStrInf_5 | VisPeri | Left | 1.35 | 3.70* |
| StriCal_1 | VisPeri | Right | 4.51*** | 3.80* |
| ExStrInf_3 | VisPeri | Left | 1.39 | 3.98* |
| ExStrInf_2 | VisPeri | Left | -0.21 | 4.17* |
| ExStrInf_4 | VisPeri | Right | 1.10 | 5.73** |
| Rsp_2 | DMNC | Left | 1.87 | 4.04* |

FDR-corrected at $q < 0.05$; *$q \leq 0.05$, **$q \leq 0.01$, ***$q \leq 0.001$

**Supplementary Table 5**. A subset of regions that demonstrate significant relationships between cortical thickness and gestation week, before (raw) and after controlling for total thickness (partial) in a multivariate regression analysis. Note that cortical thickness in regions belonging to the Visual Peripheral, Limbic A, and Default Mode C Networks becomes positively associated with gestation week after controlling for global change. This suggests that these regions decline in thickness at a rate slower than the global decrease over the gestational window. Note, this is not an exhaustive list of results; see **Supplementary File 2** for complete reporting. Abbreviations: PostC = post central; FEF = frontal eye fields; ParMed = parietal medial; Ins = insula; S2 = Somatomotor 2; PFCl = lateral prefrontal cortex; ExStrInf = extra-striate inferior; StriCal = striate calcarine; TempPole = temporal pole; Rsp = retrosplenial.

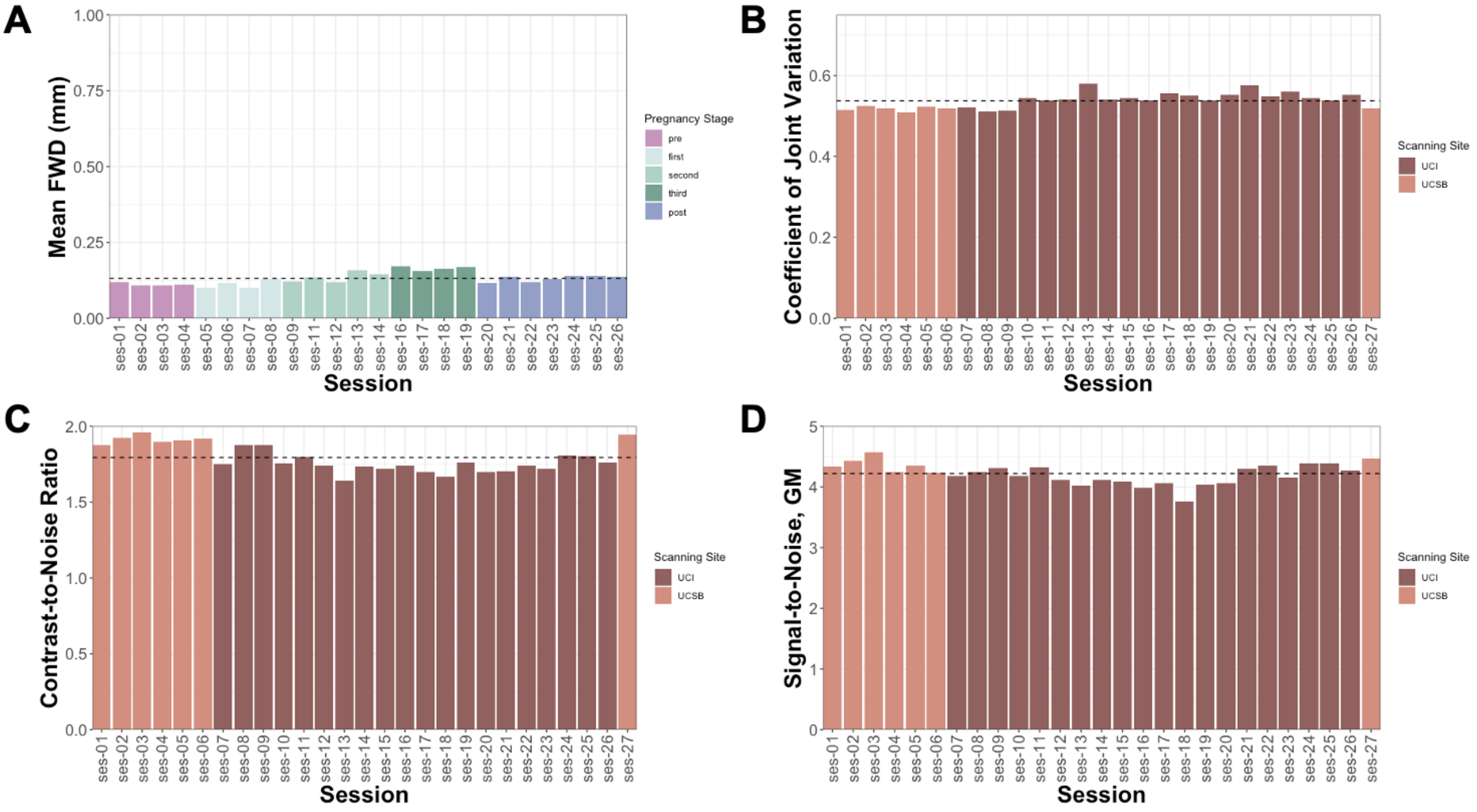


**Supplementary Figure 4.** Quality control estimates for this precision imaging experiment fell within standard ranges (see Esteban et al., 2017b); dashed horizontal line represents the average. **A)** Mean motion (i.e., framewise displacement) estimates were derived from N=17 available resting-state scans that followed each session's whole-brain T1 MPRAGE scan. Head motion was minimal compared to a conservative 'acceptable' limit of 1mm. **B–D)** Quality control estimates derived using the IQMs pipeline from *MRIQC*. **B)** Coefficient of Joint Variation is indicative of head motion and artifacts. **C)** Contrast-to-noise ratio evaluates separation of tissue distributions of GM and WM. **D)** Signal-to-noise ratio for gray matter. Cortical GMV reductions over pregnancy were still observable after controlling for these QC factors; however, the magnitude and location of these relationships were altered (see **Fig. S5**).

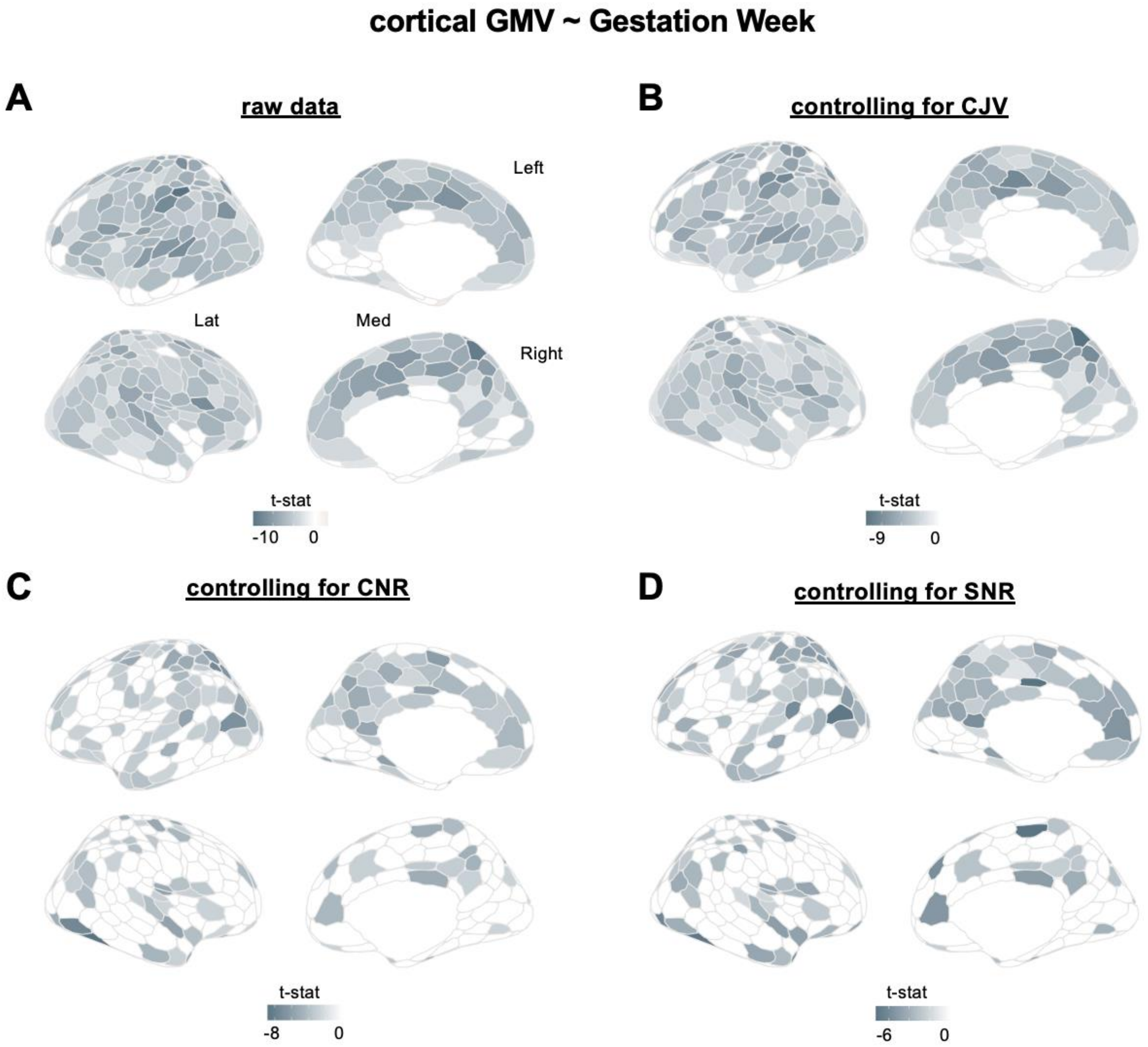


**Supplementary Figure 5.** A comparison of our raw (**A**) whole-brain GMV regression results and models (**B–C**) accounting for *MRIQC* quality control assessments (Esteban et al., 2017a). Controlling for these measures does not detract from our main finding suggesting cortical GMV reductions occur over gestation, especially within regions belonging to attention and somatosensory networks. However, there are observable decreases in CNR and SNR tied to gestation week, impacting the magnitude and location of our results. Given that data quality fell within the normal range, there may be meaningful reductions in signal to accompany volumetric reductions (e.g., regional increases in CSF paired with decreases in GMV) — a methodological nuance that warrants further exploration. *Abbreviations*: CJV, coefficient of joint variation; CNR, contrast-to-noise ratio; SNR, signal-to-noise ratio; Lat = lateral; Med = medial.

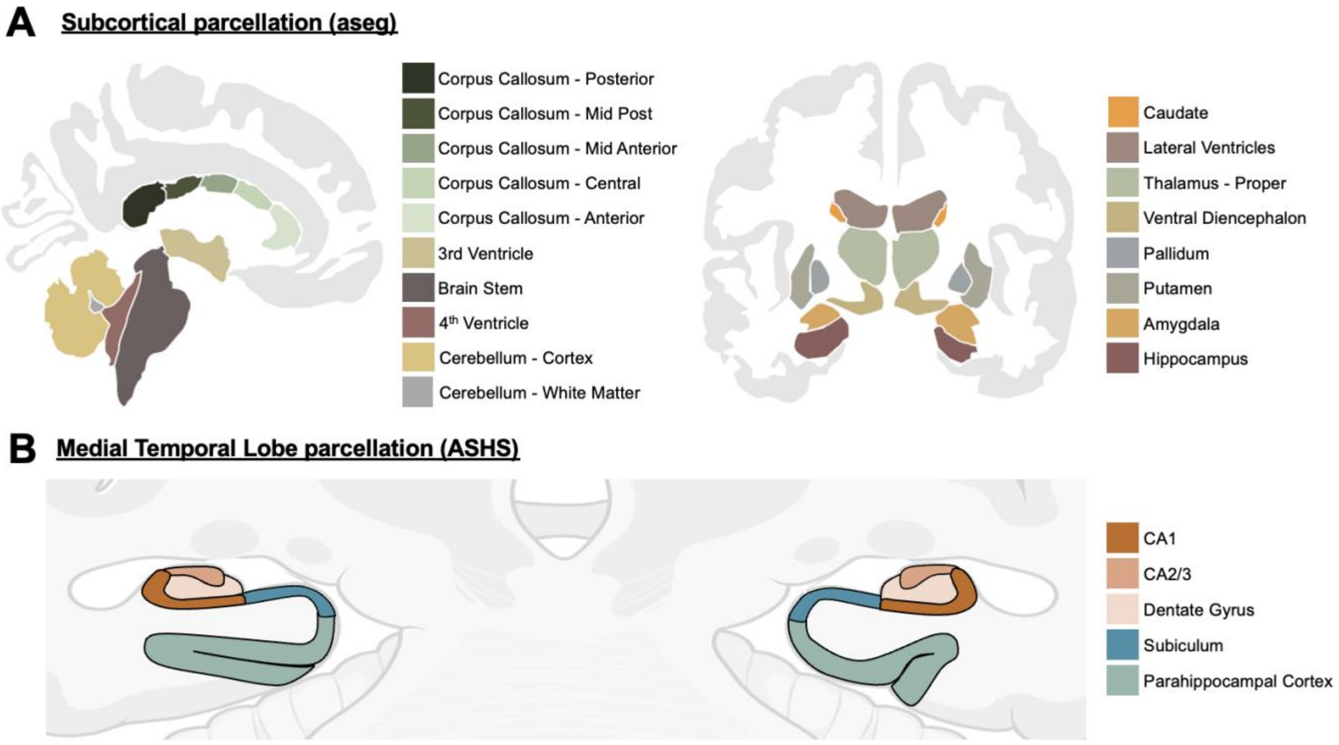


**Supplementary Figure 6.** Complete labelling of subcortical segmentations. **A)** Subcortical segmentation based upon the existence of an atlas ('aseg') containing probabilistic information on the location of structures via whole-brain T1w images. **B)** Cartoon depiction of medial temporal lobe segmentation determined via manual editing of the output of the Automatic Segmentation of Hippocampal Subfields (ASHS) software package. The participant's hippocampus and surrounding cortex were segmented into seven bilateral subregions. To note, perirhinal and entorhinal cortex subfields are not shown in this graphic due to slice coverage but are included in statistical analyses.

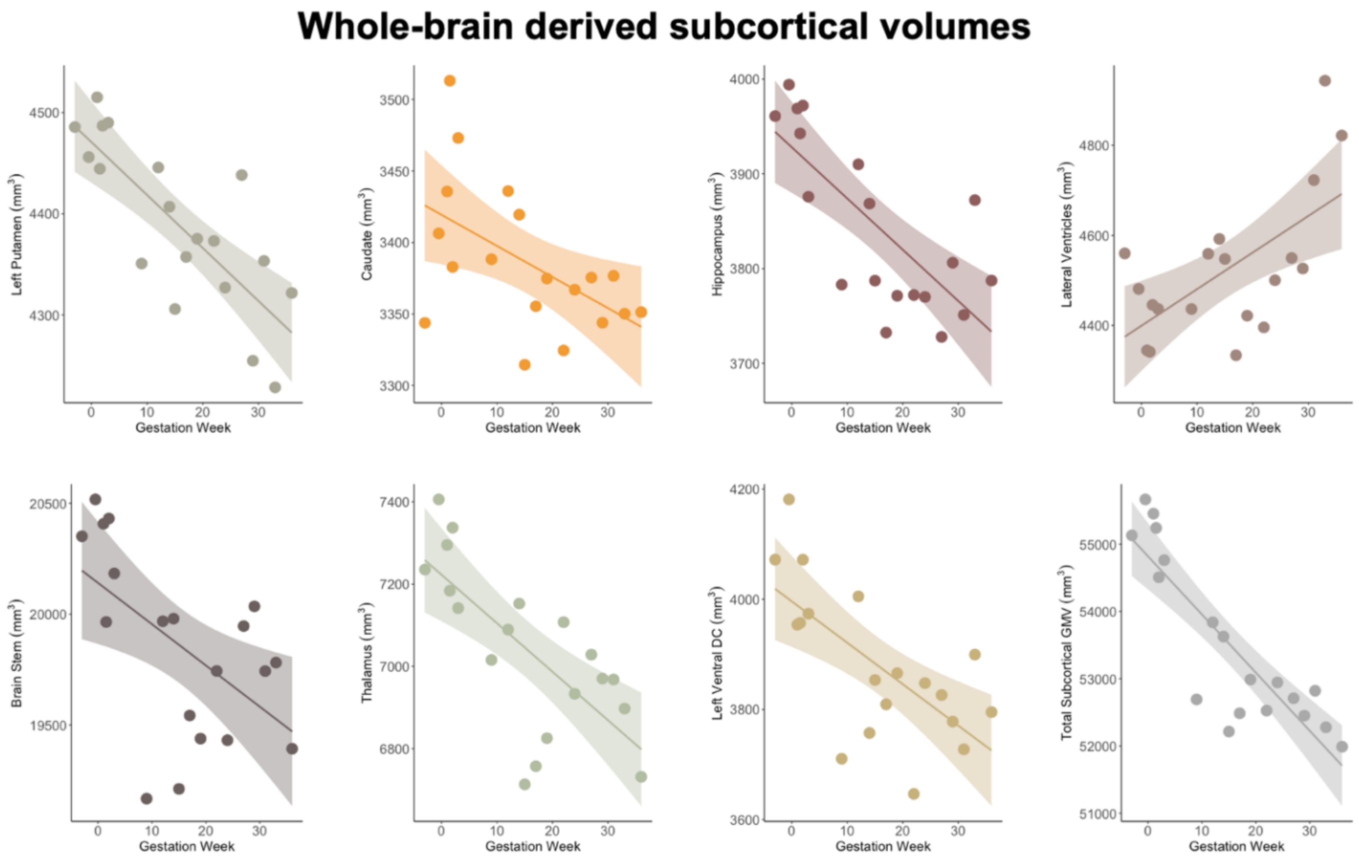


**Supplementary Figure 7.** Subcortical regions defined via the 'aseg' parcellation scheme (volume estimates derived via FreeSurfer) that demonstrate significant relationships with gestation week (baseline – 36 weeks; $q < .05$). Regions that displayed similar patterns across left and right hemispheres were averaged here; see **Fig. 3A** and **Supplementary Table 6** for details. A significant linear model was fit for each region (shaded regions represent 95% confidence interval) and data were normally distributed. However, data patterns across a handful of structures suggest the potential for non-linear trends over gestation – a possibility that should be addressed in the future with larger cohorts.

**Supplementary Table 6.** Relationships between subcortical GMV and gestation week

| aseg ROI | Hemisphere | T-stat |
|---|---|---|
| Ventral Diencephalon | Bilateral | L: -3.76**, R: -6.44*** |
| Putamen | Left | -5.39*** |
| Hippocampus | Bilateral | L: -3.35*, R: -5.10*** |
| Thalamus | Bilateral | L: -3.31*, R: -3.03* |
| Lateral Ventricle | Bilateral | L: 3.20*, R: 3.40* |
| Brain Stem | -- | -2.79* |
| Caudate | Bilateral | L: -2.60*, R: -2.68* |

FDR-corrected at $q < 0.05$; *$q \leq 0.05$, **$q \leq 0.01$, ***$q \leq 0.001$

**Supplementary Table 6**. Significant relationships between subcortical regions of interest (derived from whole-brain T1w scan using Freesurfer, see *Methods*) and gestation week over the course of pregnancy (baseline – 36 weeks).

**Supplementary Table 7.** Medial temporal subregion volumes ($mm^3$) and associations with gestation week and hormones

| | CA1 | CA2/3 | DG | Sub | ERC | PRC | PHC | Total Hipp Body |
|---|---|---|---|---|---|---|---|---|
| **Average (SD)** | **507.50 (25.45)** | **175.63 (18.60)** | 497.72 (30.12) | 265.33 (12.08) | 735.02 (44.61) | 2156.02 (67.35) | **1839.74 (76.00)** | 1445.18 (74.51) |
| **Min** | **462.34** | **144.16** | 451.15 | 244.29 | 643.16 | 2064.34 | **1684.42** | 1318.49 |
| **Max** | **554.35** | **202.48** | 561.98 | 281.68 | 836.64 | 2310.03 | **1979.07** | 1562.50 |
| **Range** | **92.01** | **58.33** | 110.83 | 37.38 | 193.48 | 245.69 | **294.64** | 244.01 |
| **Gestation Week** adjusted $R^2$ *(q)* | **0.36[a]* (0.01)** | **0.41[a]* (0.008)** | -0.06 *n.s.* | -0.06 *n.s.* | 0.03 *n.s.* | 0.11 *n.s.* | **0.58* (0.0009)** | -0.06 *n.s.* |
| **Estradiol** adjusted $R^2$ *(q)* | 0.10 *n.s.* | 0.18 *n.s.* | -0.07 *n.s.* | 0.03 *n.s.* | 0.01 *n.s.* | 0.06 *n.s.* | **0.60* (0.005)** | 0.05 *n.s.* |
| **Progesterone** adjusted $R^2$ *(q)* | -0.02 *n.s.* | 0.05 *n.s.* | -0.10 *n.s.* | -0.06 *n.s.* | -0.08 *n.s.* | 0.08 *n.s.* | 0.75 *n.s.* | -0.05 *n.s.* |

FDR-corrected at $q < 0.05$ | n.s. indicates q > 0.05 | [a] denotes a quadratic relationship, otherwise linear | * remains significant with total volume correction

**Supplementary Table 7**. Medial temporal lobe subregion volumes in relation to gestation week and sex hormones. Regression models were used to determine associations between subregion volume and gestation week, estradiol, and progesterone (baseline – 36 weeks). Non-linear (CA1, CA2/3) and linear (PHC) volumetric decreases were observable across gestation (also see **Fig. 3B**). There was no significant change in other subregions or total volume of the hippocampal body (sum of CA1, CA2/3, DG, and Sub), or in the parahippocampal gyrus (**Fig. S8**). *Abbreviations*: DG = dentate gyrus; Sub = subiculum; ERC = entorhinal cortex; PRC = perirhinal cortex; PHC = parahippocampal cortex.

### Medial Temporal Lobe Subregion Volumes

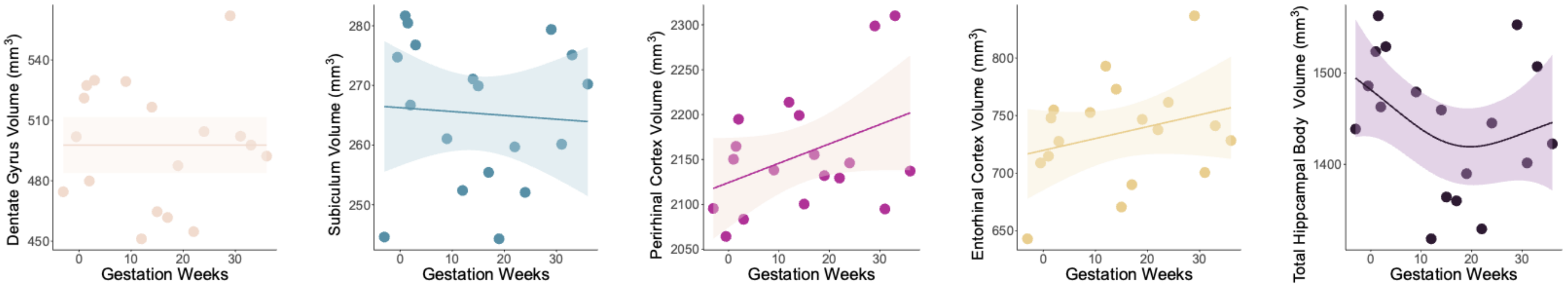


**Supplementary Figure 8.** Medial temporal subregion volumes across gestation, all non-significant at $p > 0.05$. However, data patterns across a handful of structures suggest the potential for non-linear trends over gestation – a possibility that should be addressed in the future with larger cohorts.

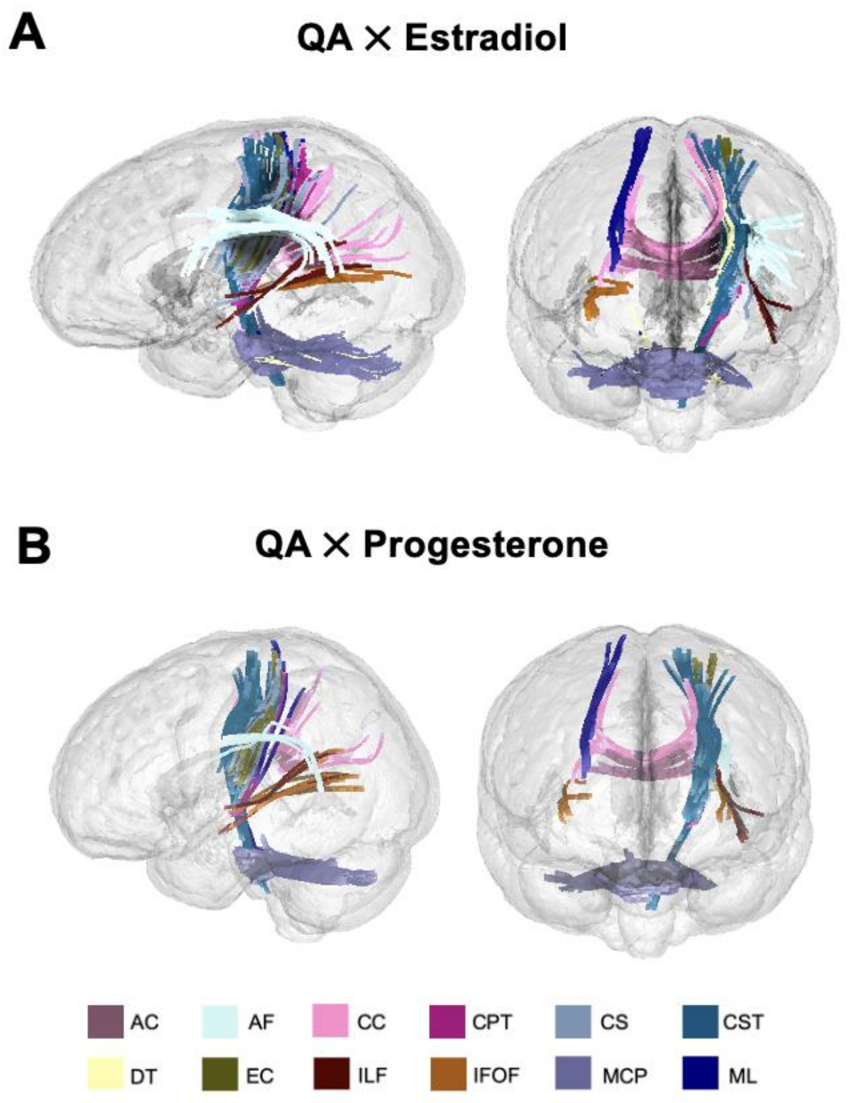


**Supplementary Figure 9.** Relationships between white matter integrity and hormones throughout gestation. **A)** White matter tracts in which quantitative anisotropy (QA) was significantly positively associated with estradiol (FDR q < .001). **B)** White matter tracts in which QA was significantly positively associated with progesterone (FDR q < .001). *Abbreviations : AC*= anterior commissure , AF = arcuate fasciculus, CC = corpus callosum, CPT =corticopontine tracts, CS = corticostriatal tracts, CST = corticospinal tracts, DT = dentothalamic tract, EC= extreme capsule, ILF = inferior longitudinal fasciculus, IFOF = inferior frontal occipital fasciculus, MCP = middle cerebellar peduncle, ML = medial lemniscus.

**Supplementary Table 9.** Full report of quantitative anisotropy results by tract that show positive associations with gestation week and sex hormones

| | Number of tracts | | | Mean length (mm) | | | Diameter (mm) | | | Volume (mm^3) | | |
|---|---|---|---|---|---|---|---|---|---|---|---|---|
| Region of Interest | *Gest Week* | *Progesterone* | *Estradiol* | *Gest Week* | *Progesterone* | *Estradiol* | *Gest Week* | *Progesterone* | *Estradiol* | *Gest Week* | *Progesterone* | *Estradiol* |
| Anterior_Commissure | 68 | – | – | 62 | – | – | 5.4 | – | – | 1412 | – | – |
| Arcuate_Fasciculus_L | 559 | 56 | 12 | 56 | 58 | 56.3 | 8.8 | 6 | 3 | 3348 | 1814 | 333 |
| Arcuate_Fasciculus_R | 215 | – | – | 60 | – | – | 6.2 | – | – | 1801 | – | – |
| Cingulum_Frontal_Parietal_L | 44 | – | – | 56 | – | – | 4 | – | – | 703 | – | – |
| Cingulum_Frontal_Parietal_R | 64 | – | – | 54 | – | – | 4.7 | – | – | 951 | – | – |
| Cingulum_Parolfactory_R | 22 | – | – | 57 | – | – | 2.1 | – | – | 204 | – | – |
| Corpus_Callosum_Body | 854 | 122 | 41 | 60 | 58 | 55.9 | 16 | 8 | 5 | 11679 | 3012 | 1075 |
| Corpus_Callosum_Forceps_Major | 2247 | 39 | 29 | 68 | 58 | 55.9 | 13 | 5 | 4 | 9513 | 1044 | 725 |
| Corpus_Callosum_Tapetum | 2367 | 49 | 33 | 60 | 57 | 54.7 | 17 | 5 | 4 | 13884 | 1144 | 762 |
| Corticopontine_Tract_Occipital_L | 37 | – | – | 55 | – | – | 3.9 | – | – | 664 | – | – |
| Corticopontine_Tract_Occipital_R | 23 | – | – | 53 | – | – | 3.7 | – | – | 558 | – | – |
| Corticopontine_Tract_Parietal_L | 58 | 55 | 12 | 53 | 66 | 68 | 5.2 | 5 | 3 | 1133 | 1330 | 557 |
| Corticopontine_Tract_Parietal_R | 77 | – | – | 57 | – | – | 5.9 | – | – | 1550 | – | – |
| Corticospinal_Tract_L | 170 | 233 | 131 | 58 | 62 | 63.1 | 8.3 | 10 | 8 | 3161 | 5118 | 3284 |
| Corticospinal_Tract_R | 80 | – | – | 59 | – | – | 5.8 | – | – | 1552 | – | – |
| Corticostriatal_Tract_Posterior_L | 239 | 16 | – | 57 | 60 | – | 8.2 | 3 | – | 2972 | 485 | |
| Corticostriatal_Tract_Posterior_R | 110 | – | – | 54 | – | – | 7 | – | – | 2089 | – | – |
| Corticostriatal_Tract_Superior_L | 11 | 102 | 35 | 51 | 58 | 57.2 | 3.1 | 8 | 5 | 395 | 2886 | 1176 |
| Dentatorubrothalamic_Tract_L | 18 | 35 | – | 57 | 70 | – | 3.5 | 6 | – | 533 | 1747 | – |
| Dentatorubrothalamic_Tract_R | 24 | – | – | 55 | – | – | 3.9 | – | – | 635 | – | – |
| Extreme_Capsule_L | 17 | 16 | 15 | 55 | 64 | 62 | 2.1 | 3 | 3 | 184 | 600 | 559 |
| Frontal_Aslant_Tract_R | 30 | – | – | 52 | – | – | 3.1 | – | – | 389 | – | – |
| Inferior_Fronto_Occipital_Fasciculus_L | 843 | – | 14 | 59 | – | 52.6 | 11 | – | 3 | 5223 | – | 391 |
| Inferior_Fronto_Occipital_Fasciculus_R | 450 | 41 | 15 | 58 | 58 | 59.3 | 9.2 | 4 | 2 | 3869 | 633 | 263 |
| Inferior_Longitudinal_Fasciculus_L | 2297 | 17 | 12 | 67 | 55 | 55.5 | 13 | 3 | 3 | 8223 | 415 | 390 |
| Inferior_Longitudinal_Fasciculus_R | 603 | – | – | 61 | – | – | 10 | – | – | 5076 | – | – |
| Medial_Lemniscus_L | – | 69 | 34 | – | 58 | 55.7 | – | 6 | 5 | – | 1666 | 984 |
| Medial_Lemniscus_R | 21 | 27 | 18 | 56 | 58 | 57.2 | 3.8 | 4 | 3 | 620 | 887 | 453 |
| Middle_Cerebellar_Peduncle | – | 593 | 239 | – | 62 | 59.9 | – | 14 | 10 | – | 9450 | 4751 |
| Middle_Longitudinal_Fasciculus_L | 161 | – | – | 53 | – | – | 6 | – | – | 1516 | – | – |
| Middle_Longitudinal_Fasciculus_R | 115 | – | – | 53 | – | – | 5.9 | – | – | 1473 | – | – |
| Superior_Longitudinal_Fasciculus1_R | 21 | – | – | 56 | – | – | 3.4 | – | – | 499 | – | – |
| Thalamic_Radiation_Posterior_L | 26 | – | – | 57 | – | – | 4.4 | – | – | 891 | – | – |
| Thalamic_Radiation_Posterior_R | 56 | – | – | 56 | – | – | 5 | – | – | 1114 | – | – |

**Supplementary Table 9**. Tract statistics for QA among tracks that were identified (via correlational tractography analysis) as being significantly positively associated with gestation week (FDR $q < 0.001$), progesterone (FDR $q < 0.001$), and estradiol (FDR $q = 0.001$). Tracts are labeled by hemisphere (L, left; R, right) and report the number of tracts, mean length, diameter, and volume. Dashes signify no significant relationships between the pregnancy outcome variable and respective tract, as determined by the FDR value determined in DSI studio.

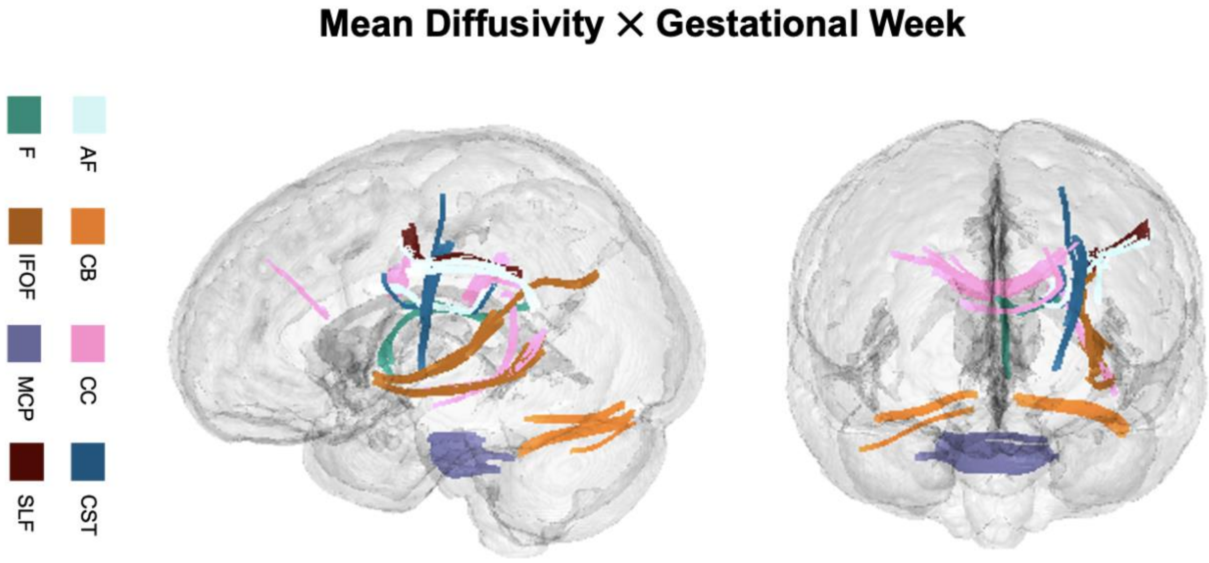


**Supplementary Figure 10.** Relationships between white matter integrity and gestation (baseline – 36 weeks). White matter tracts in which mean diffusivity (MD) was positively associated with gestational week (FDR $q < .001$). Correlations were positive, but because lower MD values typically indicate worse microstructural integrity these findings indicate tracts where microstructural integrity declined during gestation. *Abbreviations :* AF = arcuate fasciculus, CB= cerebellum, CC = corpus callosum, CST = corticospinal tracts, F= fornix, IFOF = inferior frontal occipital fasciculus, MCP = middle cerebellar peduncle, SLF = superior longitudinal fasciculus II

**Supplementary Table 10.** Full report of mean diffusivity results by tract that show positive associations with gestation week and estradiol

| | Number of Tracts | | | Mean Length | | | Diameter (mm) | | | Volume (mm^3) | | |
|---|---|---|---|---|---|---|---|---|---|---|---|---|
| Region of Interest | *Gest Week* | *Progesterone* | *Estradiol* | *Gest Week* | *Progesterone* | *Estradiol* | *Gest Week* | *Progesterone* | *Estradiol* | *Gest Week* | *Progesterone* | *Estradiol* |
| Arcuate_Fasciculus_L | 13 | - | - | 51.8 | - | - | 3.7 | - | - | 556 | - | - |
| Cerebellum_L | 27 | 18 | 29 | 55.1 | 51.6 | 52 | 3.5 | 3.3 | 4.5 | 537 | 442 | 835 |
| Cerebellum_R | 11 | 35 | 14 | 50.9 | 50.7 | 51 | 2.7 | 4.4 | 3.1 | 290 | 782 | 381 |
| Corpus_Callosum_Body | 78 | - | - | 53.4 | - | - | 6 | - | - | 1527 | - | - |
| Corpus_Callosum_Forceps_Major | - | - | 11 | | - | 53.1 | | - | 2.5 | | - | 269 |
| Corpus_Callosum_Tapetum | 21 | - | - | 53.1 | - | - | 4.2 | - | - | 743 | - | - |
| Corticospinal_Tract_L | 11 | - | - | 51.3 | - | - | 3.1 | - | - | 381 | - | - |
| Fornix_L | 11 | - | - | 52.7 | - | - | 2.5 | - | - | 255 | - | - |
| Inferior_Fronto_Occipital_Fasciculus_L | 43 | - | - | 55.3 | - | - | 4.8 | - | - | 999 | - | - |
| Middle_Cerebellar_Peduncle | 164 | 31 | 17 | 53 | 52.9 | 52 | 6.6 | 3.4 | 3 | 1801 | 488 | 357 |
| Superior_Longitudinal_Fasciculus2_L | 14 | - | - | 53.1 | - | - | 2.7 | - | - | 303 | - | - |

**Supplementary Table 10.** Mean diffusivity tract statistics for those significantly correlated to gestational week (FDR $q < 0.001$) and estradiol (FDR $q = 0.047$). DSI Studio recommends reporting tracts that with a significance level of FDR < 0.2. Based on this criteria, we have included tracts with MD associated with progesterone (FDR $q = 0.188$) for completeness. Regions are labeled by hemisphere (L, left; R, right) and reported on by number of tracts, mean length, diameter, and volume. Dashes indicate no statistical relationship. Higher MD values typically indicate worse microstructural integrity.

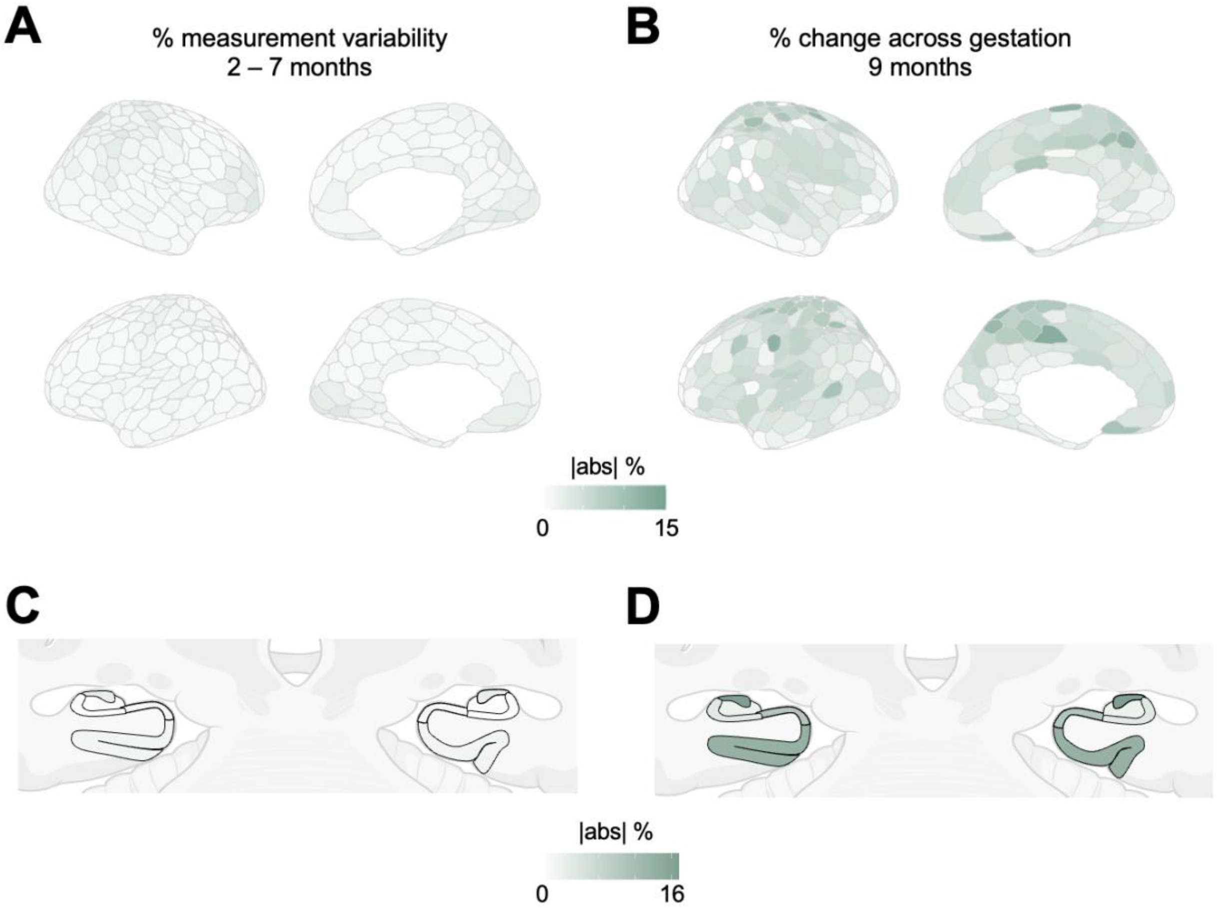


**Supplementary Figure 11.** We further contextualized our results by utilizing the Day2Day dataset from Filevich et al., 2017, wherein eight individuals (6F/2M) underwent brain imaging over an extended longitudinal window (2–7 months). Whole-brain T1w images from each subject were run through the same protocols used in our own data, ensuring proper comparison of the output from both datasets. **A)** Using Day2Day T1w data, we computed the degree of measurement variability (see *Methods*) for each region of the brain, allowing us to derive an estimate of typical day-to-day variability and measurement error. **B)** The change that occurs during pregnancy (baseline vs. 36 weeks gestation) goes far beyond what can be deemed 'typical' variability in the brain over a similar time course. A similar comparison was made for medial temporal lobe volumes which **D)** exhibit changes during pregnancy that exceed **C)** normative variability. Note: *Brain visualizations created with R-package ggseg (A-B; Mowinckel and Vidal-Piñeiro, 2020) and BioRender.com (C-D).*

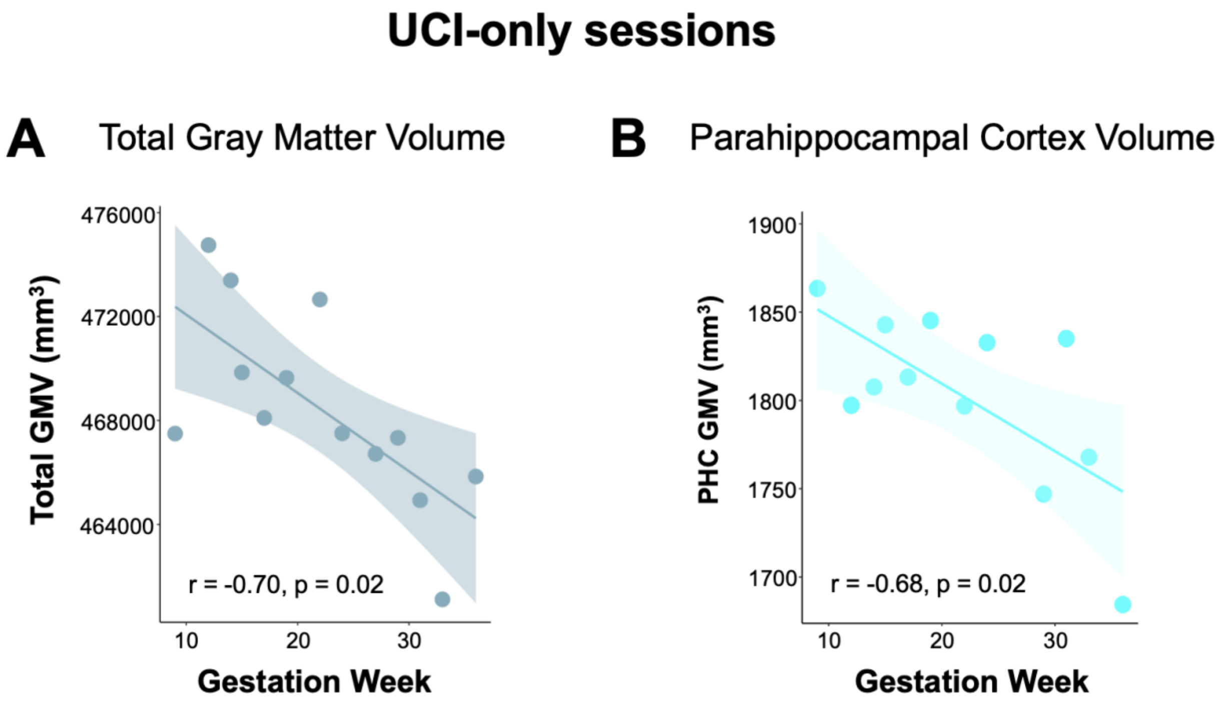


**Supplementary Figure 12.** Representative relationships between neuroanatomy and gestation when evaluating UCI sessions only (n=13). **A)** Gestation week remains significantly tied to decreasing total GMV (Pearson's product-moment correlation). Additionally, most large-scale brain networks remain negatively associated with GMV across gestation, except for Visual (Central, Peripheral), Limbic (A,B), and Default Mode Network (C) (see **Table S1** for patterns across all sessions). **B)** Gestation week remains significantly associated with decreasing parahippocampal cortex volume.
